# In vitro characterization of the baker’s yeast deubiquitinase Ubp3

**DOI:** 10.64898/2026.08.19.745719

**Authors:** Lars Bostelmann-Arp, Sakshi Khosa, Jens Reiners, Kai Mayor Völtzke, Sander H. J. Smits, Andreas S. Reichert, Lutz Schmitt

## Abstract

Ubp3 is one of about 20 deubiquitinases (DUBs) in *S. cerevisiae*. The current view generally assumes that Ubp3 requires its interaction partner Bre5, which is proposed to function as a positive regulator. Accordingly, the Ubp3/Bre5 complex has been implicated in a broad range of cellular processes for example trafficking between ER and Golgi, stress granule formation and selective autophagy. However, the molecular basis of this proposed Bre5-dependent activity remains unclear. To address this at a molecular level, Ubp3, Bre5, and related constructs were heterologously expressed in *E. coli*, purified to homogeneity, and characterized *in vitro*. Both proteins contain folded domains as well as extensive intrinsically disordered regions (IDRs). Despite this structural complexity, the Ubp3/Bre5 complex could be isolated following either co-expression *in vivo* or after *in vitro* assembly. Unexpectedly, complex formation with Bre5 was not required for the catalytic activity of full length Ubp3. Furthermore, even the isolated catalytic domain of Ubp3 was fully active against two distinct substrates in the absence of Bre5, demonstrating that its deubiquitinating activity is intrinsically independent of Bre5. These findings indicate that the catalytic domain alone is sufficient for substrate cleavage, whereas the extensive IDRs of Ubp3 and its cofactor Bre5 might contribute to substrate recognition or specificity. Overall, this study challenges the prevailing model of Bre5-dependent activation of Ubp3 and provides new insights into the molecular organization of the Ubp3/Bre5 system. More broadly, it highlights the importance of intrinsically disordered regions in regulating deubiquitinase function and cellular signaling networks.

## Introduction

Ubiquitin is a 76 amino acids small protein present in every eukaryotic cell and its reversible conjugation through its C-terminal glycine to substrate proteins serves as an important post translational modification. It is attached through a three-step enzymatic cascade via ubiquitin-activating (E1), ubiquitin-conjugating (E2), and ubiquitin-ligating enzymes (E3) [1, 2]. In most cases ubiquitin is attached through an isopeptide bond to the ⍰-amino group of a lysine residue displayed on the surface of the target protein. As ubiquitin itself has seven lysins and its N-terminus, it can accept additional ubiquitin modifications on eight different sites. Proteins can be mono-ubiquitinated, multi-mono-ubiquitinated as well as polyubiquitinated. The latter can form either linear or branched ubiquitin chains. On top of this, ubiquitin linkages through other amino acids than lysine and even non-protein targets such as sugars have been described in recent years [1]. These various ubiquitin modifications, the so-called ‘ubiquitin code’ [2], adopt distinct topologies and lead to different cellular outcomes. These include among many others the tagging of proteins for degradation through the proteasome, the ubiquitin-proteasome pathway (UPS), but have also been shown to change activity and localization of proteins and to regulate numerous cellular pathways, for example quality control, ribophagy and mitophagy, respectively [3]. More recently, also the critical role of a C-terminally extended isoform of ubiquitin, termed CxUb, which needs to be matured and conjugated via the ‘ubiquitin fusion degradation’ pathway in cellular quality control pathways, such as mitophagy, and ageing was reported [4].

These examples stress the importance of ubiquitin modifications in multiple cellular processes and emphasize that a tightly regulated balance between ubiquitination and the reversal of this process, deubiquitination, is essential for cellular homeostasis and survival. Key to this balance are deubiquitinases (DUBs) that can cleave or modify ubiquitin moieties. They are also required for the upkeep of the free ubiquitin pool by either recycling chains from the UPS or through cleavage of newly translated ubiquitin fusion proteins. In humans, there are approximately 100 DUBs, grouped into 7 families [5], while in yeast 22 DUBs belonging to five of these different classes have been described [6].

The largest DUB family in yeast and in humans contains the ubiquitin-specific proteases [5, 6]. They are characterized by a conserved fold that is flanked by varying C- and N-terminal extensions. This so-called USP-fold has been first described by Hu *et al.* [7] and is comprised of an architecture described as an open hand with finger, palm and thumb subdomains. This architecture forms a specific S1 site, which binds the distal ubiquitin and locates its C-terminus to the catalytic triad. Often times this consists of Cys, His and Asp/Asn amino acids [7]. These domains are structurally highly conserved but their sequence can vary. A study by Ye *et al.* also divided this fold into six different segments which comprise the binding site. The connecting loops serve as places for specific domain insertions such as Zn^2+^-binding domain [8].

The yeast deubiquitinase ubiquitin carboxy-terminal hydrolase 3 (Ubp3), which is homologue to human USP10, forms a complex with brefeldin-A sensitivity protein 5 (Bre5). Based on previous studies, Bre5 has been proposed to function as a positive regulator of Ubp3 and has been suggested to be required for its activity [9]. Accordingly, the Ubp3–Bre5 complex has been implicated in regulating ER-to-Golgi transport through modulation of the ubiquitination of Sec23 and βP-COP [9, 10]. Another study proposed that this also influences temporal regulation affecting stringent quality control. [11].

A number of additional functions have been assigned to Ubp3 or the Ubp3/Bre5 complex, often using knockout strains. For example, Ubp3 deubiquitinates RNApolII [12], which requires Bre5 binding to RNA [13]. Ubp3 knockout reduces proteasomal degradation, which in turn increases proteotoxicity [14]. The formation of stress granules requires Bre5 and active Ubp3 [15] and relies upon the intrinsically disordered region (IDR) of Ubp3 [16]. On the other hand, only limited structural data are available for these proteins. Two studies by Li *et al.* revealed that both proteins form a hetero-tetrameric complex and interact through a binding motif in the N-terminus of Ubp3 and the nuclear transport factor 2 (NTF2)-like domain of Bre5 [17, 18].

In a genome wide screen by Müller *et al*. [19], Ubp3 and Bre5 were identified as negative regulators of mitophagy and it was shown that the complex dynamically relocates to mitochondria upon rapamycin treatment. In contrast, other types of selective autophagy such as cytosol to vacuole trafficking [20] or ribophagy [21, 22] are positively regulated by the complex. During ribophagy, Ubp3 is specifically required for degradation of the ribosomal core particle, but not for the regulatory particle [23]. This process further depends on interactions with Cdc48 and is antagonized by the E3 ubiquitin ligase Rsp5 [24]. Additionally, the absence of the Cdc48 cofactor Ubp3 results in mitochondrial aggregation and fragmentation [25].

Although many of these functions have been inferred from genetic studies, several putative Ubp3 substrates have been identified, for example Sec23 and β’-cop [9, 10], Atg19 [20], Tbp1 [26] and eS7A [27]. In addition to these ubiquitinated proteins, CxUb was identified as a substrate of Ubp3, which has a role in stress resistance, mitophagy and longevity [4].

Despite the wealth of information about the various functions of the complex, little is known about their molecular mode of action and the substrate recognition. Here we report a detailed *in vitro* characterization of Ubp3 and Bre5. We demonstrate that both proteins contain large IDRs, which complicated their biochemical characterization. Nevertheless, the Ubp3/Bre5 complex could be readily isolated by either co-expression of both proteins or assembly from isolated proteins. Most notably, our analysis of catalytic activity challenges the prevailing model that Bre5 acts as a positive regulator and that complex formation is required for Ubp3 activity [9], Instead, the isolated catalytic domain of Ubp3 is fully competent to cleave multiple substrates, even in the absence of its N-terminal region and Bre5. These findings demonstrate that the intrinsic catalytic activity of Ubp3 is independent of Bre5 *in vitro*.

## Results

In order to enable the biochemical and biophysical characterization of the proteins investigated in this study, recombinant expression and purification protocols were established for all constructs. Details of the production of proteins for subsequent *in vitro* characterization are provided in Materials and Methods. Proteins were routinely purified through consecutive purification steps using affinity and gel filtration columns (Table S2). All proteins eluted in a symmetric or nearly symmetric peak by size exclusion chromatography (SEC) (Figure 1A-C). SDS-PAGE and Western Blot verified the presence of Bre5 and Ubp3 in the respective peaks (Figure 1D-F). It is important to stress that purification of full-length Ubp3 was only possible if tags at both termini were used, as the protein was prone to proteolytic degradation. Bre5 only showed a single band at 70 kDa while Ubp3 and its catalytic domain (Ubp3^Cat^, residues 407-912) displayed major bands at 100 kDa and 60 kDa respectively. Both, Ubp3 and Ubp3^Cat^, also showed additional lower molecular weight bands compared with the full-length proteins. These were confirmed by Western blot as degradation products, as they contained only one or the other of the two terminal affinity tags (Figure 1D,E).

**Figure 1:**
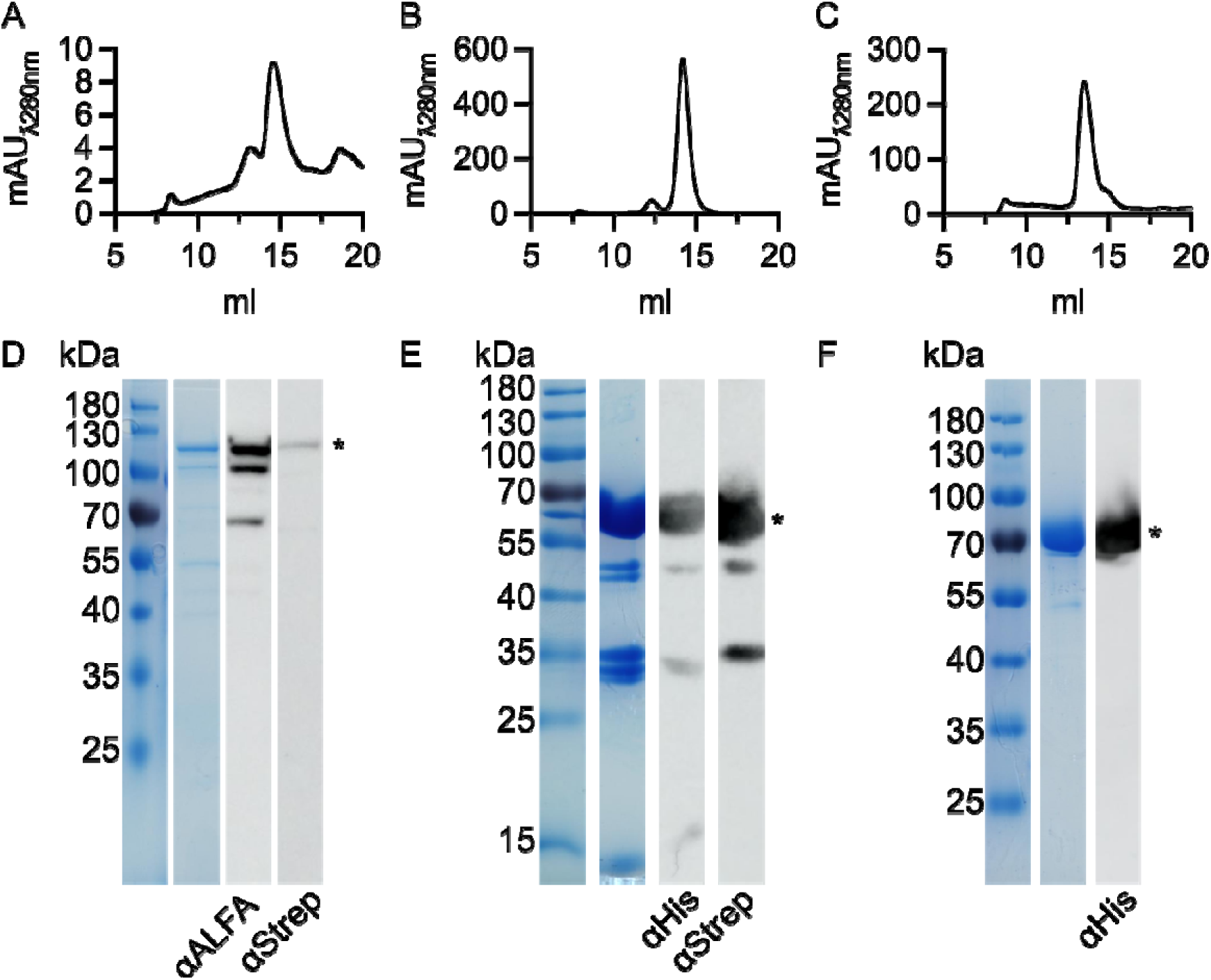
Purification of the individual constructs. Gel filtration chromatograms of Ubp3 eluting at 15 ml (A), its catalytic domain Ubp3^Cat^ eluting at 14.5 ml (B) and Bre5 eluting at 14 ml (C). Corresponding SDS-PAGE with marker proteins and Western-Blot for Ubp3 (D), its catalytic domain (E) and Bre5 (F), asterisks mark the respective band of the full-length proteins.

To investigate the stability and overall oligomeric state of the purified proteins, they were analyzed using NanoDSF [28] and Size-Exclusion Chromatography coupled with Small-Angle X-ray Scattering (SEC-SAXS) [29, 30]. For the Bre5 dimer, AlphaFold3 [31] predicted that only the nuclear transport factor 2 (NTF2)-like domain and the RNA recognition motif (RRM) would adopt a tertiary structure, while the rest would remain in the form of a random coil. This was confirmed by SEC-SAXS, which determined that Bre5 forms a dimer in solution and exists as a flexible and elongated particle based on the *p(r)* function and Kratky plot (see Figure S3 and Table S4). We used the created AlphaFold3 model as a starting template, keeping the NTF2-like dimer interface intact and treating the RRM domain as a flexible domain. We applied an Ensemble Optimization Method (EOM) analysis to obtain information about the flexible behavior of the Bre5 dimer in solution. The EOM analysis revealed a broad, elongated conformational distribution (Figure S3, Rg and Dmax distribution of Bre5) rather than compact orientations, which aligns well with the experimental data (χ^2^-value = 1.1) (Table S4). An example model from the EOM analysis to visualize the flexibility of the Bre5 dimer is shown in Figure 2B (the whole ensemble can be found in the supplementary information Figure S4). In addition, the NanoDSF measurements showed an initially high ratio (Supplementary Figure S1) and only a slight minimum in the first order derivative (Figure 2A), which could be the result of unfolding in the RRM and NTF2-like domains, that would be reflected through changes in tyrosine fluorescence properties. There were no classic transitions from low to high signal ratios, suggesting that the tryptophans of Bre5 were already solvent exposed. Therefore, the NanoDSF data further supports the SAXS model of a flexible mostly disordered protein.

For Ubp3 and the catalytic domain of Ubp3 (amino acid residues 407-912; Ubp3^Cat^), the NanoDSF signals followed the classical unfolding behavior transitioning from a low to high ratio (Supplementary Figure S1), which in turn leads to a maximum in the first order derivative (Figure 2C). For both proteins the melting temperature (T_M_) was almost identical (Ubp3: 47.82 ± 0.03 °C, Ubp3^Cat^: 47.09 ± 0.16 °C), despite the full-length protein containing a larger number of tryptophane residues. This suggests that the catalytic domain of Ubp3 is the only folded part of the protein, while the N-terminal part is intrinsically disordered. The SEC-SAXS analysis confirmed that Ubp3 is a monomer in solution (Supplementary Table S4). Structural characterization of the data, via the *p(r)* function and the Kratky plot, revealed that Ubp3 is an elongated molecule with a folded core in solution. Within the EOM analysis, Ubp3 exhibited a bimodal-like behavior in solution, existing in compact (about 33 %) or super-elongated state (two models with 22 and 44 % respectively) (Supplementary Figure S5, RG dmax) (Figures 2D). With a χ^2^-value of 0.959, the ensemble is in great agreement with the experimental data. This behavior is driven by the flexible N-terminal tail of Ubp3. On the other hand, Ubp3^Cat^ is almost completely structured (Figure 4A). Based on an AlphaFold3 model and a created AFflecto library [32], only a few flexible loops underwent modification. This was based on the scoring of the models against the experimental SEC-SAXS data (Supplementary Figure S8, Table S4) for the description of the in-solution behavior of the Ubp3^Cat^ core part. Overall, these results demonstrate that both Bre5 and Ubp3 contain large intrinsically disordered regions (IDRs), with only some domains adopting classical folds in solution.

**Figure 2:**
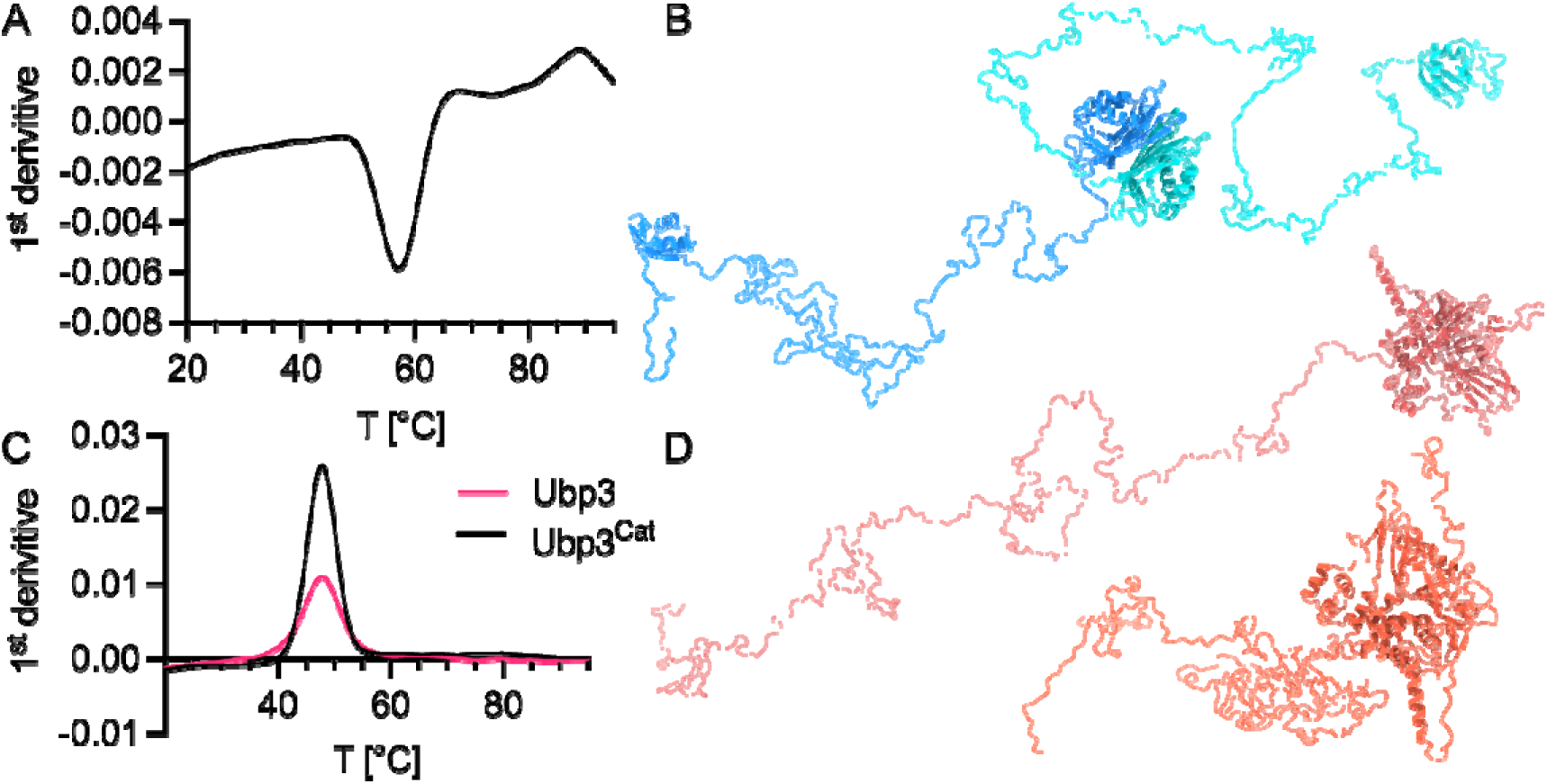
A: Analysis of Bre5 and Ubp3. First derivative of the unfolding of Bre5, **B:** one representative EOM model generated from the SEC-SAXS data for Bre5 with one monomer in cyan and one in blue, folded domains are in cartoon representation. **C**: First derivative of the unfolding curves for Ubp3 (red) and Ubp3^Cat^ (black). **D**: Two representative EOM models for Ubp3 with the catalytic domain in cartoon representation and the flexible N-terminus as ribbon. One model with a more elongated state of the N-terminal IDR (upper, dark red) and one model in the more compact state (lower, bright red) (SEC-SAXS models not with the same scale. Further models and data can be found in the supplementary part).

We next investigated, whether the previously described Ubp3-Bre5 complex could be reconstituted *in vivo* as well as *in vitro* and which stoichiometry it would adopt, as there had been conflicting results [17]. To investigate *in vivo* complex formation, inactive Ubp3 (Ubp3^C469A^), Bre5 and ubiquitin were co-expressed from a single plasmid and subsequently purified. To achieve the purification two affinity purifications were performed, where the first step utilized the tag on Bre5 and the second step the tag on Ubp3, before finally isolating the complex via size-exclusion chromatography (SEC). The resulting chromatogram (Figure 3A) showed the presence of multiple species. However, all proteins were detected by Western blot analysis in the peak eluting at 13 ml (Figure 3B). Other SEC peaks contained individual proteins, for example, Ubp3 eluted at 15 ml. The eluting fraction of the assembled complex was concentrated and used for SEC-SAXS. The collected data revealed a large, elongated, multidomain molecule with a certain degree of flexibility, as demonstrated by the p(r) function and the Kratky plot (Figure S9, Table S4), and a molecular weight of 347.39 kDa was determined using MoW2 [33] analysis. This indicated a heterohexamer (molecular weight of 348.33 kDa) consisting of two Bre5 molecules, two Ubp3 molecules and two ubiquitin molecules (Table S4). We created an AlphFold3 model of this complex and compared it with the experimental data. Due to the large IDR regions of the complex, the resulting model showed a high degree of mismatch with the experimental data (χ^2^ value of 64.86), particularly in the low s-region (<1 nm⁻¹). To improve agreement with the experimental data, we used BilboMD [34], keeping the NTF2-like dimer interface and the Ubp3 binding interface intact, while treating the RRM and Catalytic domains as flexible. The experimental data were best approximated by a three-state model, in which the complex existed either as a fairly compact state (about 8 %) (Figure 3E) or as a more elongated state (two states of about 42 and 48 % respectively), with the NTF2-like domains mostly in the vicinity of Ubp3^Cat^ and large flexible loops (Figure 3D). Overall, modelling proved to be difficult due to the large IDRs in both proteins. In summary, the complex relies on Bre5 dimerization, to which Ubp3 is tethered via its binding domain. These results demonstrate that the proteins form a stable complex *in vivo*. As further validation of functionality of our *in vitro* purified proteins, we analyzed the potential of complex assembly by incubating the individual components together and isolating the complex by SEC. Therefore, individual proteins were first analyzed on a microÄkta (GE / Cytiva). The peaks corresponding to the proteins (Figure 3C, Ubp3 red line, Bre5 cyan line) were concentrated and mixed in a 1:1 molar ratio. This resulted in one single peak at earlier elution volume (Figure 3C black line), which indicates assembly of a larger complex even in the absence of ubiquitin. The elution volume of 1.3 ml in a 10 times smaller column volume also matches that of the *in vivo* complex, eluting at 13 ml.

Furthermore, we investigated whether ubiquitin can bind to the catalytic domain alone, without the flexible N-terminal region. In the SEC-SAXS experiment, Ubp3^Cat^ in complex with ubiquitin eluted as a clear peak, with the excess of ubiquitin eluting later (Figure S11). Analysis of the SAXS data revealed a 1:1 ratio of Ubp3^Cat^ to ubiquitin. The resulting *p(r)* function and the Kratky plot revealed that, upon substrate binding, the catalytic domain remains folded in a globular state with minor elongation, likely due to the loop regions. To analyze the in-solution structural behavior of this complex, we created an AlphaFold3 docking model of the Ubp3^Cat^/ubiquitin complex. Based on our knowledge of the isolated catalytic domain, we created a library of different loop positions in the docking model. We then scored the resulting library against the experimental data to find the best-fit model (Figure S11, Table S4), which had a χ^2^ value of 1.019. The model showed ubiquitin nested in the catalytic domain, with its C-terminal glycine residues located at the active site, giving an overall buried surface area of 2,112 Å^2^ as calculated in ChimeraX [35]. In comparison to the crystal structure of Usp7 with ubiquitin aldehyde (PDB code: 1NBF) [7] (Figure 4B), ubiquitin is positioned identically in the Ubp3^Cat^ model and the Usp7 structure. More importantly, the overall fold of the two USP domains aligns well, with only three loop regions in Ubp3^Cat^ being larger and finding no match in the crystal structure. Overall, the proteins do form a complex, regardless of the conditions as a heterotetramer of two Bre5 molecules and two Ubp3 molecules, and each Ubp3 molecule can bind a ubiquitin molecule through its catalytic domain in a 1:1 stoichiometry.

**Figure 3:**
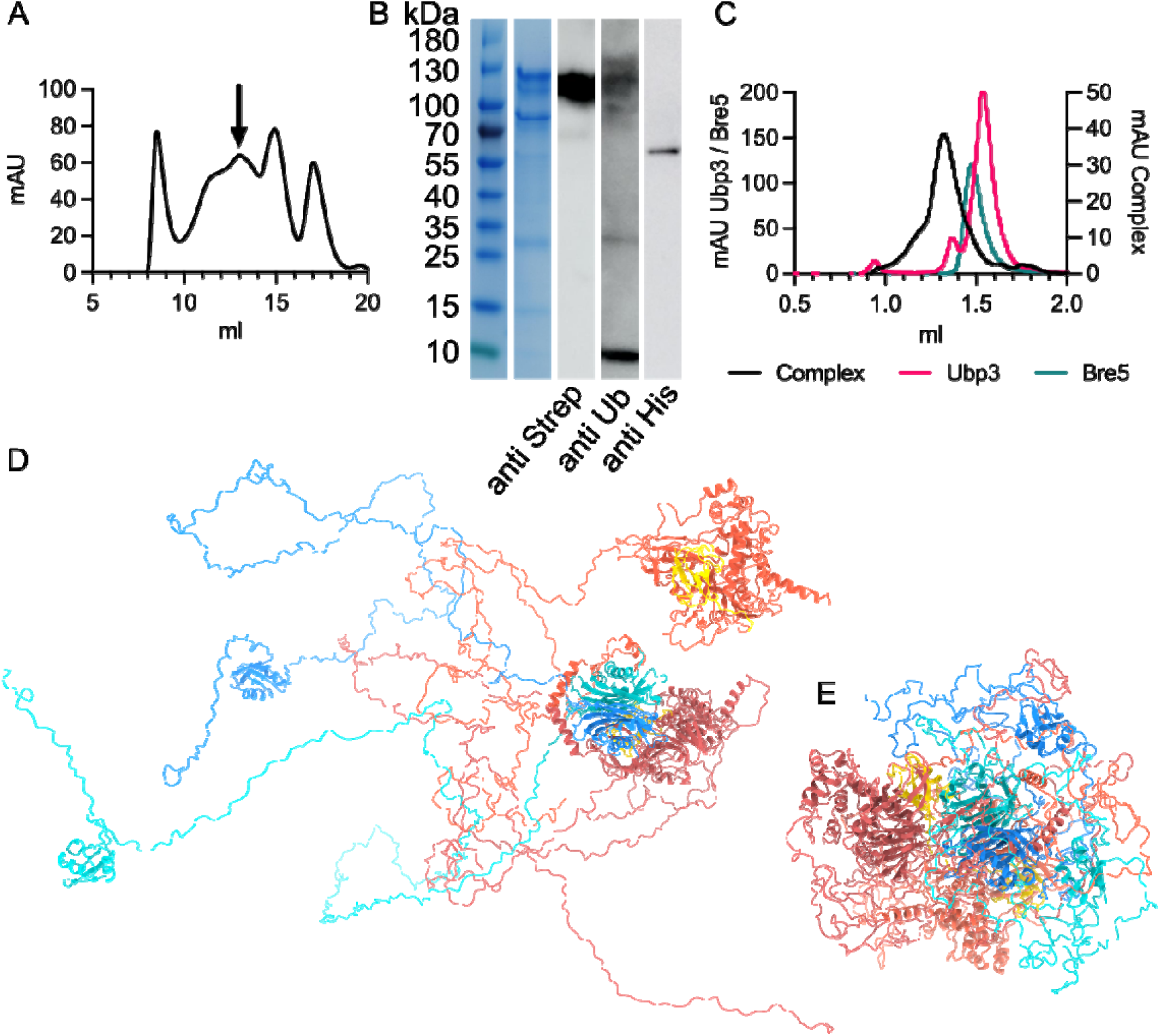
Analysis of the Ubp3-Bre5 complex. **A**: SEC of the Ubp3-Bre5-Ub complex formed *in vivo* with the arrow marking the complex. **B**: SDS-PAGE gel and Western blot of the full complex formed *in vivo*. **C**: SEC of the individually purified proteins on a microÄKTA (GE / Cytiva) with Ubp3 in red, Bre5 in cyan and the complex in black. *D:* One of two elongated flexible models that are part of the three-state SAXS model for the complex *E:* Third model of the three-state model, showing the complex in a more compact conformation, Ubp3 in red (dimers in different shades of red), Bre5 in blue (dimer shown in different shades of blue and ubiquitin in yellow (Models D and E with different scales. Further models and data can be found in the supplementary part).

In order to see whether either of the observed interactions induce additional folding or lead to stabilization, we tested unfolding with NanoDSF [36]. The complex of Ubp3 and Bre5 did only show a melting temperature of 47.19 ± 0.01 °C (supplementary data Figure S1), which matched that of the catalytic domain. This indicates that the Bre5-Ubp3 complex is either not stable and dissociates at higher temperatures or that there is no folding of the IDRs into a tertiary structure upon complex formation, which could shield the tryptophan residues. When the catalytic domain was tested together with a two-fold molar excess of ubiquitin, no change in the T_M_ was observed (Figure 4C). Therefore, the catalytic domain does bind ubiquitin as seen in SEC-SAXS but this interaction is not sufficient to induce any further stabilization of the catalytic domain. These results indicate that neither of the interactions underlying formation of the different complexes induce major folding events nor increase the stability of the catalytic domain.

**Figure 4:**
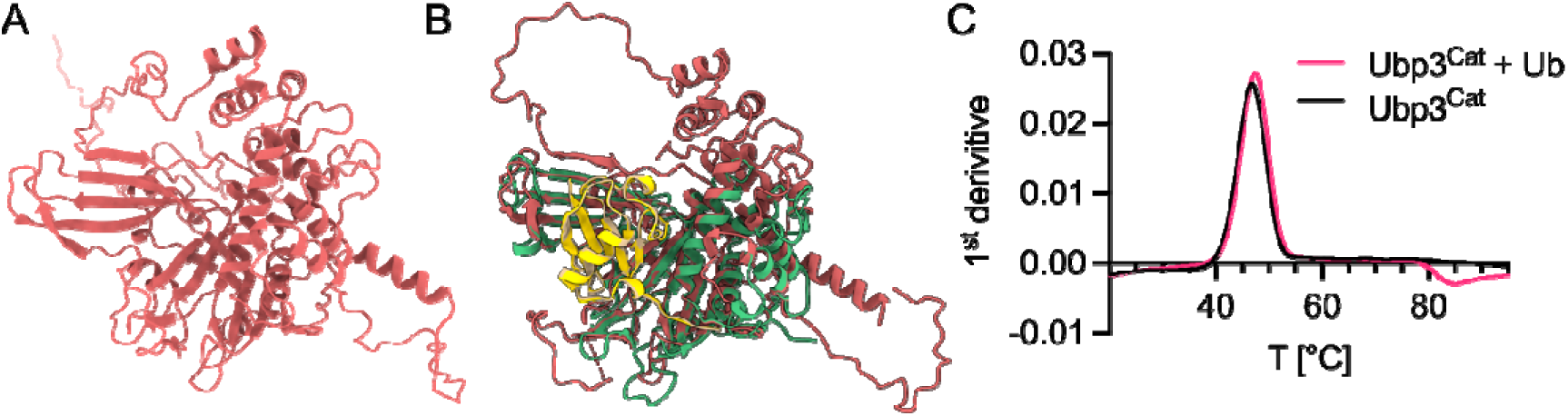
A: Analysis of Ubp3^cat^. Aflecto model for the Ubp3 as determined by SEC-SAXS. **B**: Superimposition of the Usp7-Ub-aldehyde structure (green and orange) PDB: 1NBF with the Ubp3^Cat^-Ub model derived from SEC-SAXS (red and yellow respectively). **C**: NanoDSF comparison of Ubp3^Cat^ in the presence or absence of ubiquitin.

Having established the mode of complex formation and its interaction with ubiquitin, we next asked how this affects the activity of Ubp3. To investigate this, we used three different substrates to confirm the activity of the proteins or their complexes *in vitro*. First, we used the classical Ub-AMC activity assay [37], that has been employed in the characterization of several other DUBs for example [38–40]. It relies on the change in fluorescence properties of 7-amido-4-methylcoumarin (AMC) if it is cleaved from the C-terminus of ubiquitin. We measured fluorescence changes over time for Ubp3, Bre5, the positive control Usp2, which was supplied by the manufacturer (Cayman chemicals) and the catalytic inactive variant of Ubp3, namely Ubp3^C469A^ [9] (Figure 5A). Bre5, and the inactive mutant showed no increase in fluorescence over time, indicating no release of AMC and therefore no activity. In contrast, both the positive control and Ubp3, showed a clear increase in fluorescence over time, indicating that these proteins can process Ub-AMC and release AMC. This is in agreement with a previous study [24], but raises questions regarding the functional role of Bre5 in Ubp3-mediated deubiquitination. Bre5 had been previously postulated to act as a positive regulator of Ubp3 activity [9]. Since we have demonstrated that Bre5 and Ubp3 form a complex from their individual components, we also investigated whether different amounts of Bre5, and therefore altered stoichiometries of the individual components could modulate Ubp3 activity. Using the Ub-AMC assay, we could not detect any changes in the initial fluorescence increase of Ubp3 alone or in combination with Bre5 at molar excess ranging from 0.5 to 10-fold (Figure 5B). Thus, in our hands Ubp3 does not require its proposed positive regulator Bre5 to cleave Ub-AMC *in vitro* and activity is not affected by the presence of Bre5. To see whether this holds true for a substrate with an actual protein fused to ubiquitin, instead of the rather small AMC, the second substrate that we analyzed was ubiquitin fused to the stable GFP (Ub-GFP), which we purified from *E. coli* ΔelaD [41] (further details in Material and Methods). After purification, Ub-GFP was incubated in presence or absence of purified Ubp3 and samples were taken after 0, 10 and 20 minutes and analyzed by SDS-PAGE. Here, a single continuous protein band at 40 kDa corresponding to Ub-GFP was observed in absence of Ubp3 (Figure 5C). However, in the presence of Ubp3, the signal intensity of these bands decreased over time and a second signal at approximately 30 kDa, corresponding to GFP without attached ubiquitin, appeared and its signal intensity increases over time. These results show that Ubp3 is also capable of cleaving a Ub-GFP fusion protein without the requirement of Bre5. Knowing that the isolated catalytic domain is sufficient for ubiquitin binding, we also tested whether the catalytic domain alone was sufficient for activity using again Ub-AMC. Unexpectedly, the isolated catalytic domain also efficiently cleaved Ub-AMC (Supplementary Figure S2). To not only see activity as fluorescence increase but also determine kinetic parameters, we resorted to a third substrate. This was CxUb from *S. cerevisiae*, ubiquitin extended with a C-terminal asparagine, termed Ub^77N^ in this study. This had been recently described as a substrate of DUBs, including Ubp3 [4]. Ub^77N^ was expressed and purified as described in Materials and Methods and incubated with Ubp3, the Ubp3-Bre5 complex or Ubp3^Cat^ for 15 min at different substrate concentrations (Figure 5D). This enabled determination of the Michaelis-Menten kinetics as summarized in Table 1. As observe before, Ubp3 and the Ubp3-Bre5 complex showed similar behavior with comparable biochemical parameters. Ubp3^Cat^ on the other hand showed a higher K_M_, as well as a higher V_max_. Summarizing these different activity assays, *in vitro* Bre5 does not appear to act as a positive regulator for deubiquitination itself, and importantly Ubp3^Cat^ alone is sufficient for catalytic activity as demonstrated by the catalytic efficiency k_cat_/K_M_ (Table 1), which is identical within experimental error for Ubp3^Cat^, Ubp3, and the Ubp3-Bre5 complex.

**Table 1:** Kinetic parameters of Ubp3, the Ubp3-Bre5 complex and the catalytic domain, as determined with Ub^77N^. Parameters calculated from three measurements with SE.

| Protein | $K_M$ ($\mu M$) | $v_{max}$ ($\mu mol * min^{-1} * l^{-1}$) | $k_{cat}$ ($s^{-1}$) | $k_{cat}/K_M$ ($s^{-1} * M^{-1}$) |
| --- | --- | --- | --- | --- |
| Ubp3 | $176.8 \pm 31.5$ | $5.2 \pm 0.3$ | $0.43 \pm 0.03$ | $2443 \pm 450$ |
| Ubp3-Bre5 | $131.1 \pm 28.3$ | $4.2 \pm 0.3$ | $0.35 \pm 0.02$ | $2675 \pm 652$ |
| Ubp3<sup>Cat</sup> | $302.5 \pm 20.6$ | $9.0 \pm 0.2$ | $0.75 \pm 0.02$ | $2471 \pm 176$ |

**Figure 5:**
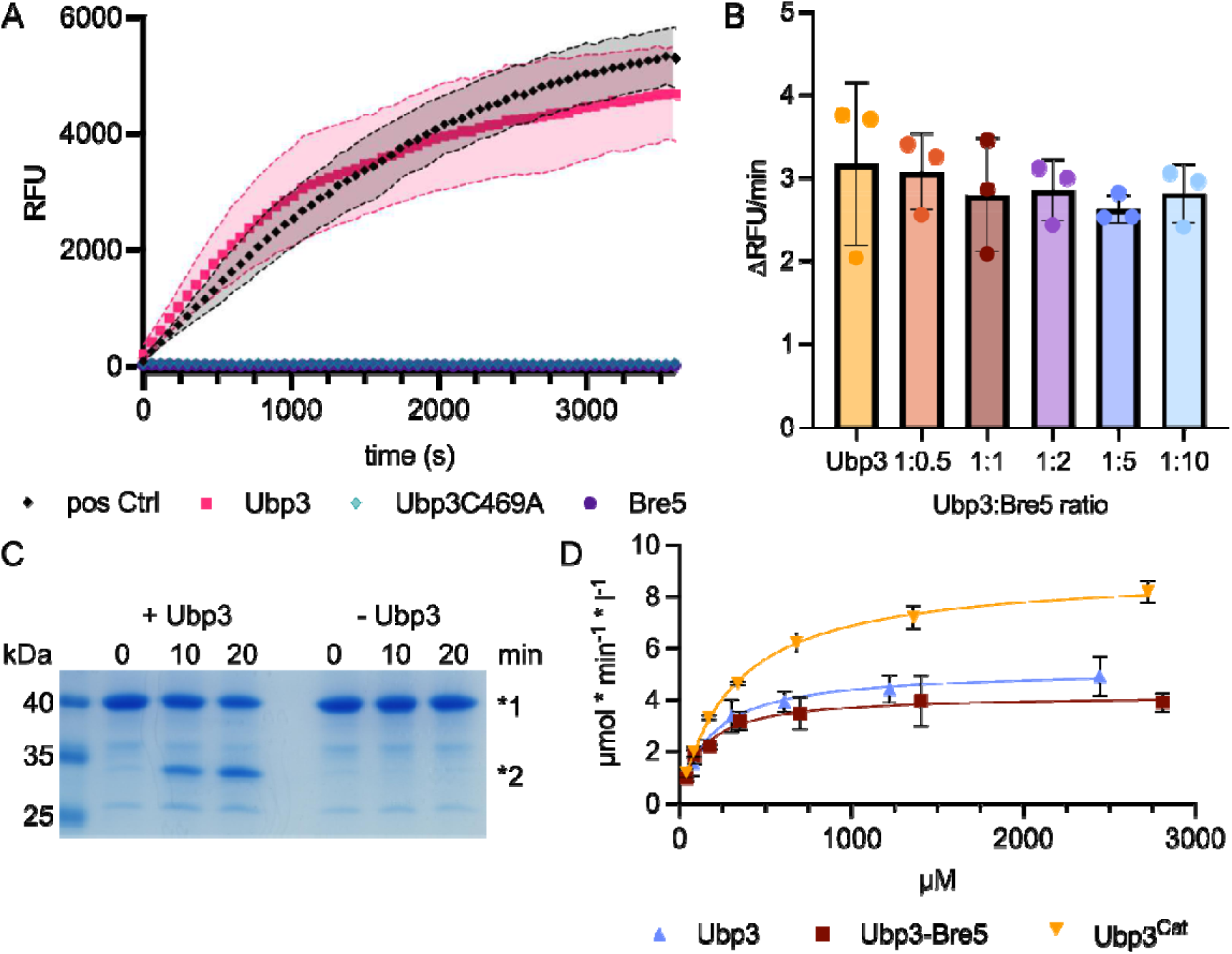
A: Kinetic analysis of Ubp3. Ub-AMC cleavage assay, fluorescence increase over time in RFU for the positive control (black diamonds), Ubp3 (red squares), Ubp3C469A (cyan diamond) and Bre5 (purple spheres), shaded areas show the SD of three replicates. **B**: Rate of initial fluorescence increase in ΔRFU/min for Ubp3 and samples with different ratios of Ubp3:Bre5 in the Ub-AMC assay. **C**: SDS-PAGE showing the cleavage of Ub-GFP over time with (+) or without (-) Ubp3. Molecular weight standards are shown to the left. *1 indicates the signal for Ub-GFP and *2 indicates the cleaved GFP. **D**: Reaction velocity (µmol * min^-1^ * l^-1^) of Ubp3-(red squares), Ubp3-Bre5 complex-(yellow inverted triangles) and Ubp3^Cat^-(blue triangles) mediated cleavage of Ub^77N^ at different substrate concentrations (µM). (Lines in the same color show the respective Michaelis-Menten equation fitted. Error bars represent the SD of 3 replicates).

## Discussion

In this study, we described the successful isolation of Ubp3, its catalytic domain and Bre5. SEC-SAXS and NanoDSF demonstrated that both proteins contain large IDRs, while the catalytic domain of Ubp3 adopts a folded state, that fits the canonical USP fold [8] and binds ubiquitin with a 1:1 stoichiometry. Ubp3, Bre5 and ubiquitin form a complex with a 2:2:2 stoichiometry when co-expressed *in vivo*, which can also be assembled from the individual components *in vitro*. Most important, we demonstrated that the catalytic activity is already present in the isolated catalytic domain of Ubp3 and neither Bre5 nor the N-terminal IDR of Ubp3 are required.

In contrast to several *in vivo* studies involving Ubp3 knockouts [14, 21, 23], we focused on the *in vitro* characterization based on a detailed heterologous expression approach. Although Ubp3 was previously purified from *E. coli* [9, 17], we optimized the expression and purification strategy to obtain homogeneous protein in sufficient yields for our subsequent studies. However, the purification process already hinted at the disordered nature of the proteins, as degradation products of Ubp3 were consistently observed [42] and the yields of Ubp3 and Bre5 were reduced due to aggregation and liquid-liquid phase separation (LLPS), a common characteristic of intrinsically disordered proteins [43]. On the other hand, the catalytic domain showed only limited degradation and could be purified with high yields. Additionally, the melting temperature (T_M_) observed in NanoDSF measurements of Ubp3 and its catalytic domain was identical. This implies that only this catalytic domain undergoes unfolding, thereby showing that it is folded in the first place.

While Li *et al* [17] reported the crystal structure of a complex of the Bre5 NTF2-like domain and a Ubp3-derived peptide, we explored the structure of the full-length proteins. Using SEC-SAXS, we determined an ensemble of models that featured some folded domains and large flexible IDRs. This architecture for yeast DUBs has also been observed for Ubp15 [44]. Using this method, we could also verify that Ubp3 alone, and for that matter also its catalytic domain, remain monomeric, while Bre5 forms a dimer. Bozza *et al* [44] showed that the short stretches of the Ubp15 IDR interact with specific substrates. The presence of the IDR in both our proteins suggests that this could also be the mode of substrate recognition for Ubp3. In this context, Bre5 could be a cofactor that extends the range of substrates which would fit to the plethora of pathways in which the complex is involved. For example, the RRM of Bre5 also extends the activity to RNA-related targets [13]. The IDRs are also relevant for the stress granule formation as described [16], and as commonly observed with the Bre5 homologue G3bp1 which has been shown to undergo LLPS [45]. Within the SAXS analysis we observed mostly elongated, flexible models. However, a small percentage (about 33 % for Ubp3 and about 8 % for the complex) existed in a fairly compact state. This compact state could explain how a substrate, after recognition through the IDRs, could be oriented in a way that allows a bound ubiquitin to enter the binding pocket within the catalytic domain of Ubp3.

The complex assembled *in vitro* and the complex isolated following *in vivo* co-expression showed similar behavior in SEC. This indicated a similar hydrodynamic radius, which was not visibly altered by the presence or absence of ubiquitin, probably due to its small size and masking effects caused by the IDRs. The SEC-SAXS analysis of the complex formed *in vivo* showed a heterohexamer with a 2:2 stoichiometry for Ubp3 and Bre5. This is in line with the complex architecture previously observed by Li *et al.* [17]. While some deubiquitinases can undergo autoinhibition upon oligomerization [46], we did not observe reduced activity of Ubp3 in the presence of Bre5 in the Ub-AMC assay. These findings suggest that in the case of Ubp3 Bre5-mediated oligomerization serves a different role.

In contrast to the generally accepted proposal that Bre5 acts as a positive regulator of Ubp3 [9], our *in vitro* results challenge this view. We demonstrated that both Ub-AMC and Ub-GFP were readily processed by Ubp3 in the absence of Bre5. Furthermore, Ub-AMC cleavage occurred with comparable initial velocities regardless of the amount of Bre5 present. Importantly, the catalytic domain alone was active against Ub-AMC and furthermore, also active against a recently described substrate Ub^77N^ (CxUb) [4]. As we could only observe a linear relation between initial reaction velocity and substrate concentrations for feasible, low Ub-AMC concentrations, we performed kinetic analysis using Ub^77N^. By varying the substrate concentration, we determined K_M_ and V_max_ values for Ubp3, the Ubp3-Bre5 complex and the catalytic domain of Ubp3. The observed differences between Ubp3 or Ubp3-Bre5 and the Ubp3^Cat^ could be explained by two factors or a combination of both. In the assay, frozen protein was used, and as Ubp3 was less stable compared to the catalytic domain alone, the actual concentration of Ubp3 in the assay could be artificially lower due to aggregation and degradation. On the other hand, the difference would also make sense, as the catalytic domain is smaller and therefore more motile and it also has fewer flexible domains which might hinder accessibility of the ubiquitin binding site. Both of these factors could allow for a higher turnover, as seen in the experimental data. Compared to other USPs, the K_M_ value of Ubp3 is approximately 100-fold higher [40]. One possible explanation for such a discrepancy could be the presence of two loops that might obstruct the binding site, as described for USP14 [47]. In the case of Ubp3, these loops are even more extensive. In general, the binding of ubiquitinated substrates to the USP fold and the release of cleavage products should also be considered. The buried surface area of 2112 Å^2^ between ubiquitin and Ubp3 is quite large and therefore the binding and release kinetics could impact the overall reaction velocity and not just the cleavage reaction itself. Lastly, this value is also considerably higher than the natural physiological ubiquitin concentration of approximately 85 µM [48]. Taken together, these observations suggest that substrate recognition and proper localization might be much more important than the cleavage reaction, which would make sense for an enzyme involved in multiple different pathways.

In summary, we show that Ubp3 is an intrinsically active deubiquitinase whose catalytic domain is sufficient for substrate cleavage *in vitro*. Although Ubp3 interacts with Bre5 through its intrinsically disordered N-terminal region, this interaction is not required for its intrinsic catalytic activity. Instead, our findings support a model in which Bre5 and the non-catalytic regions of Ubp3 might contribute to substrate recognition, subcellular localization, and stability of Ubp3.

## Materials and Methods

### Plasmids and strains

For heterologous expression, genes from *S. cerevisiae* (kindly provided on plasmids from ASR) were cloned into pQLink plasmids using Gibson assembly [49], as per the manufacturer’s protocol (NEB). Mutations and truncations were done using site directed mutagenesis. Co-expression plasmids were created using the method described for pQLink plasmids [50]. A detailed overview of the different constructs and their sequences can be found in the supplementary material. The plasmids were transformed into *E. coli* BL21(DE3), ΔmalP or ΔelaD strain, according to Table S2. For expression, 2 l LB medium supplemented with 100 mg/ml ampicillin were inoculated to an OD_600_ of 0.1 using an overnight culture. Cells were first grown to OD_600_ 0.3 – 0.4 at 37 °C with shaking at 180 rpm, then cooled to 18°C and grown further at 18 °C, 180 rpm until OD_600_ 0.6 – 0.8 was reached, upon which the expression was induced with 1 mM IPTG. Cells were further grown at 18 °C, 180 rpm for the time indicated in Table S2 and harvested by centrifugation at 5000 g for 15 min. The supernatant was discarded and cell pellets were stored at – 20 °C until further usage.

### Purification of proteins for *in vitro* analysis

Cell pellets were thawed on ice and resuspended in the respective resuspension buffer supplemented with 10 µg/ml of RNAse and DNAse, and 1 mM MgCl_2_. For all constructs except the His-Ubp3^Cat^-TS, SH-Ubp3^Cat^, Ub^77N^ and Ub a protease inhibitor tablet (cOmplete™, Roche, Germany) was added. The cells were passed once through a Constant Systems CF1 cell disrupter at 1.55 kbar and the lysate was clarified by centrifuging at 200,000 g for 1 h at 4 °C. Afterwards all purification steps were done at 4 °C and samples were kept on ice in between.

ALFA-Ubp3-TS and ALFA-Ubp3^C469A^-TS were resuspended in buffer A (20 mM Tris pH 7.5, 500 mM NaCl) and purified by loading the lysate onto a StrepTrap XT column (Cytiva) equilibrated with buffer A. Bound protein was washed using the same buffer and eluted using the buffer A supplemented with 50 mM biotin. Elution fractions were passed twice over ALFA selector CE resin (NanoTag Biotechnologies) equilibrated with buffer A. The protein was eluted with 500 µM ALFA elution peptide. Eluted protein fractions were concentrated using an Amicon® Ultra Centrifugal Filter (Millipore). The concentrated protein was further purified by size exclusion chromatography using a Superose 6 increase column (Cytiva) equilibrated with buffer A.

His-Ubp3^Cat^-TS was resuspended in buffer B (50 mM Tris pH 8, 50 mM NaCl, 10 % glycerol) and the lysate was adjusted to 20 mM imidazole before being loaded onto a HiTrap™ IMAC HP column (Cytiva) pre-equilibrated with buffer C (20 mM Tris pH 8, 250 mM NaCl, 20 mM imidazole). The protein was washed with 75 mM imidazole and eluted with 500 mM imidazole in buffer C. Elution fractions were loaded on a StrepTrap XT column equilibrated with buffer D (100 mM Tris pH 8, 150 mM NaCl, 1 mM EDTA). The protein was eluted with 50 mM biotin and the fractions were concentrated using an Amicon® Ultra Centrifugal Filter (Millipore). The concentrated protein was subjected to size exclusion chromatography using a Superdex 200 pg column (Cytiva) pre-equilibrated with buffer E (50 mM Tris pH 8, 150 mM NaCl).

SH-Ubp3^Cat^ was purified as described above for His-Ubp3^Cat^-TS. After the second affinity purification step, the tags were removed using NumaCut (Numaferm, Germany) at a concentration of 100 U per 0.75 mg of protein in a reaction buffer according to the manufacturer’s protocol. The reaction was incubated at 30 °C for 3 h and the target protein was collected as flowthrough using reverse IMAC with the initial conditions. The protein was further subjected to SEC as described above using a Superdex 200 increase column (Cytiva).

Bre5-His was resuspended in buffer B and purified using a HiTrap™ Chelating HP column (Cytiva) pre-equilibrated with buffer F (20 mM Tris pH 8, 250 mM NaCl). The column was washed and impurities were removed with 10 mM histidine in buffer F before eluting the protein with 100 mM histidine. Elution fractions were diluted to 50 mM NaCl using 20 mM Tris pH 7.5 and loaded on a HiTrap™ SP HP column (Cytiva) equilibrated with buffer G (50 mM NaCl, 20 mM Tris pH 7.5). The column was first washed with buffer G followed by 250 mM NaCl, and the protein was eluted using a gradient from 250 to 550 mM NaCl. The second peak containing the purest Bre5-His was concentrated using an Amicon® Ultra Centrifugal Filter (Millipore) and subjected to size exclusion chromatography using a Superose 6 column (Cytiva) and buffer H (50 mM Tris pH 7.5, 150 mM NaCl).

Ub-GFP-TS was resuspended in buffer B and purified by loading the supernatant onto a StrepTrap XT column (Cytiva) equilibrated with buffer H. The protein was eluted with 50 mM biotin and concentrated using an Amicon® Ultra Centrifugal Filter (Millipore). The concentrated protein was subjected to size exclusion chromatography using a Superdex 200 increase column (Cytiva) and buffer H. S-Ub^77N^ was resuspended in buffer B and purified similarly to Ub-GFP-TS, except for using buffer D. Following affinity purification, the protein was subjected to size-exclusion chromatography on a Superdex 75 pg column (Cytiva) equilibrated with buffer E.

Ubiquitin (Ub) was resuspended in 50 mM NaAc pH 5 and purified by heating the clarified lysate to 85 °C and incubation for 5 min under gentle stirring. Heat-induced aggregates were removed by centrifugation at 100,000 × *g* for 1 h at 4 °C. The supernatant was diluted four-fold with 50 mM NaAC pH 5 and loaded on a HiTrap SP FF column (Cytiva), equilibrated with the NaAc. Ub was eluted using a NaCl gradient and concentrated using an Amicon® Ultra Centrifugal Filter (Millipore) before subjected to size exclusion chromatography using a Superdex 75 pg column (Cytiva) and 50 mM Tris pH 7.5.

The Full complex of Ubp3^C469A^-TS, Bre5-His and Ub^3G^ was resuspended in buffer B and purified by loading the lysate, adjusted to 20 mM imidazole, onto a HiTrap™ IMAC HP column (Cytiva) equilibrated with buffer I (20 mM Tris pH 7.5, 500 mM NaCl, 20 mM imidazole). The protein was washed and Impurities were removed by increasing the imidazole concentration to 50 mM before eluting the protein with 500 mM imidazole in buffer I. Elution fractions were loaded on a StrepTrap XT column (Cytiva) equilibrated with buffer J (20 mM Tris pH 7.5, 500 mM NaCl) and washed. The protein was eluted using buffer J supplemented with 50 mM biotin. Elution fractions were concentrated using an Amicon® Ultra Centrifugal Filter (Millipore) and subjected to size exclusion chromatography using a Superose 6 increase column (Cytiva) with buffer J.

After SEC all proteins were further concentrated with Amicon® Ultra Centrifugal Filters (Millipore). Concentrations were determined on a NANODROP ND-1000 using extinction coefficients as determined from sequence (Table S1 in Supplementary information) and proteins were flash frozen in liquid nitrogen and stored at – 80 °C until use.

Protein purity was assessed by SDS–PAGE followed by staining with SERVA Quick Coomassie Stain. Protein identity was confirmed by Western blot analysis using the primary antibodies listed in Table S3. Immunoblots were developed using WESTAR ETA C ULTRA 2.0 chemiluminescent substrate according to the manufacturer’s instructions. Gels and Western blots were imaged using an Amersham ImageQuant 800.

### Ubp3-Bre5 Complex formation

Purified ALFA-Ubp3-TS and Bre5-His were thawed on ice and then separately injected onto a Superose 6 increase 3.2/300 column (Cytiva) at 4 °C using buffer H (50 mM Tris pH 7.5, 150 mM NaCl). Peak fractions of the respective peaks were concentrated using Amicon® Ultra Centrifugal Filter (Millipore). The concentrated proteins were mixed in a 1:1 molar ratio and incubated on ice for 5 min, and subsequently loaded onto the same column as described for the individual proteins.

### NanoDSF experiments

NanoDSF measurements were done using the Nano Temper Prometheus NT.Plex operated with PR.ThermControl v2.1.2 software. All measurements were carried out in duplicate using Prometheus standard capillary chips. Bre5-His, ALFA-Ubp3-TS and the corresponding complex (prepared by mixing equal volumes of the two proteins prior to measurement) were analyzed at a final respective concentration of 0.75 mg/ml and an intensity of 50 %. The catalytic domain SH-Ubp3^Cat^, with the tags removed, was measured at a concentration of 11.25 mg/ml, both alone and in presence of twofold molar excess of ubiquitin, at an intensity of 40 %.

### SAXS-Experiments

The SEC-SAXS data of His-Ubp3^Cat^-TS, SH-Ubp3^Cat^ with Ub and Bre5-His was collected on the P12 beamline (PETRA III, DESY Hamburg [51]). The P12 beamline was equipped with a PILATUS 6M detector (Dectris) at a fixed distance of 3.0 m. The measurements were performed at 20°C with a protein concentration of 12.00 mg/ml for His-Ubp3^Cat^-TS, 5 mg/ml (∼90 µM) SH-Ubp3^Cat^ (tags removed) + 200 µM ubiquitin for the complex and 6.6 mg/ml for Bre5-His, respectively. The SEC-SAXS runs were performed on Superdex200 increase 10/300 GL or Superose6 increase 10/300 GL columns (Cytiva) (100 µl inject), with a flowrate of 0.6 ml/min. 2400 frames were collected for each protein sample with an exposer time of 0.995 sec/frame. Data were collected and scaled to absolute intensity against water.

The SEC-SAXS data of ALFA-Ubp3-TS and the Bre5-Ubp3-ubiquitin complex (full complex) was collected on beamline BM29 at the ESRF Grenoble [52]. The BM29 beamline was equipped with a PILATUS 2M detector (Dectris) at a fixed distance of 2.827 m. The measurements were performed at 20°C with a protein concentration of 2.15 mg/ml for ALFA-Ubp3-TS and 7.44 mg/ml for the Bre5-Ubp3-ubiquitin complex. The SEC-SAXS runs were performed on a Superos6 increase 10/300 GL column (Cytiva) (100 µl inject) at a flowrate of 0.6 ml/min. 1200 frames were collected with an exposer time of 2 sec/frame. Data were scaled to absolute intensity against water.

All used programs for data processing were part of the ATSAS Software package (Version 3.0.5) [53]. Primary data reduction was performed with the programs CHROMIXS [54] and PRIMUS [55]. The Guinier approximation [56] was used to determine the forward scattering *I(0)* and the radius of gyration (*R*_g_). The pair-distribution function *p(r)* was created with the program GNOM [57] and determined the maximum particle dimension (*D*_max_).

AlphaFold3 [31] was used to predict initial models of His-Ubp3^Cat^-TS. SH-Ubp3^Cat^ with ubiquitin in complex, ALFA-Ubp3-TS, Bre5-His and the Bre5-Ubp3-ubiquitin complex.

AFflecto [32] and respectively, RANCH (part of Ensemble Optimization Method (EOM)) [58, 59] were used to create an ensemble of models with remodeled flexible parts. The initial AlphaFold3 model was compared with CRYSOL [60] and scored the created library to find the best fit model within the ensemble. For multistate ensembles the full EOM pipeline [58, 59], or BilboMD [34] were used while keeping docking interfaces and domains intact.

### Activity measurements of Ubp3 and Ubp3^Cat^

Ub-AMC assays were performed using 10 nM purified ALFA-Ubp3-TS, ALFA-Ubp3^C469A^-TS or Bre5-His and 5 µM Ub-AMC. Samples were prepared in assay buffer (50 mM HEPES pH 7.5 at 20 °C, 0.5 mM EDTA, 0.1 mg/ml BSA, 1 mM DTT) in a 96-well plate with a final reaction volume of 50 µl. Samples were equilibrated at room temperature for 20 min before addition of the substrate. Fluorescence increase was monitored for 1 h using a TECAN Infinite M Plex plate reader with i-control 2.0 software, with excitation and emission wavelengths set to 360 nm and 460 nm, respectively. For the measurement with varying Ubp3-Bre5 ratios, 10 nM Ubp3 was mixed with the respective amount of Bre5 and incubated for 20 min prior to substrate addition. The reaction was then monitored for 1 h under the same conditions. Initial velocities were determined as slope between 1 and 9 min.

The kinetic analysis of His-Ubp3^Cat^-TS, ALFA-Ubp3-TS and ALFA-Ubp3-TS with Bre5-His was performed with S-Ub^77N^ as substrate in 40 µl reaction volume. A concentration series of S-Ub^77N^ was prepared in assay buffer (50 mM Tris pH 7.5 at 20 °C, 150 mM NaCl, 5 mM DTT, 0.05 % Tween 20) and reactions were initiated by adding the respective enzymes. Enzymes were prepared in assay buffer (ALFA-Ubp3-TS and Bre5-His were mixed and incubated on ice for 30 min prior to starting the reactions) with a final concentration of 200 nM in the assay. The reaction was incubated for 15 min at 20 °C and terminated by heating the samples at 95°C for 5 min. Precipitated protein was pelleted at 20238 g for 5 min. Subsequently, 30 µl of the supernatant was added to 950 µl of 0.05 % ninhydrin in ethanol and incubated at 37 °C for 4.5 h. The samples were centrifuged again and the supernatant was analyzed on a Varian Cary 50 scan photometer over a wavelength of 300-650 nm. Absorbance values at 356 nm were converted to asparagine concentrations using an asparagine standard prepared in assay buffer and treated identically to the cleavage reactions. Michaelis-Menten fit was done using GraphPad Prism version (11.0.2).

Purified Ub-GFP was incubated with or without ALFA-Ubp3-TS in SEC buffer (50 mM Tris pH 7.5 at 4 °C, 150 mM NaCl) at 20 °C. Samples were collected after 0, 10 and 20 min and the reaction was terminated by addition of SDS-loading dye (100 mM Tris ph 6.8 at 20 °C, 4 % SDS, 0.02 % bromphenolblue, 40 % glycerol, 5 mM DTT), followed by heating at 95 °C for 5 min

## Supporting information

Supplementary Information with 4 tables and 12 figures

## Abbreviations

IDR: Intrinsically disordered region
DUB: deubiquitinase
Ub: Ubiquitin
SEC: Size-Exclusion Chromatography
SEC-SAXS: SEC-Small-Angle X-ray Scattering
NanoDSF: Nano differential scanning fluorimetry
EOM: Ensemble Optimization Method
AMC: 7-Amino-4-methylcoumarin
GFP: Green fluorescent protein
RFU: Relative fluorescence unit
LLPS: liquid liquid phase separation
UPS: Ubiquitin proteasome pathway
CxUb: C-terminally extended ubiquitin
NTF2: Nuklear transport factor 2
RRM: RNA recognition motif

## Data availability

The SAXS data have been deposited to the Small Angle Scattering Biological Data Bank [61] with the accession codes XXX. Collected SAXS raw data from the ESRF (proposal ID’s MX-2485, MX-2603 and MX-2696) can be found under DOI XXX.

## Acknowledgments

We acknowledge DESY (Hamburg, Germany), a member of the Helmholtz Association HGF, for the provision of experimental facilities. Parts of this research were carried out at PETRA III and we would like to thank Clement Blanchet, Cy M. Jeffries, Dmytro Soloviov and Timur Tropin (EMBL Hamburg) for their assistance in using beamline P12.

We acknowledge the European Synchrotron Radiation Facility (Grenoble, France) for provision of synchrotron radiation facilities under proposal ID’s MX-2485, MX-2603 and MX-2696 and we would like to thank Petra Pernot, Mark Tully and Dihia Moussaoui for their assistance in using beamline BM29.

