## Supplementary Information with 4 tables and 12 figures for "In vitro characterization of the baker’s yeast deubiquitinase Ubp3"

**Constructs**

Ubp3 (ALFA-Ubp3-Trs-TS)

Ubp3^C469A^ (ALFA-Ubp3(C469A)-Trs-TS)

Ubp3^Cat^ (6xHis-Tev-Ubp3(407-912)-Trs-TS)

Ubp3^Cat^-Ub-Complex (S-His-Tev-Ubp3(407-912) + Ub) (Tags removed for experiments)

Bre5 (Bre5-Trs-10xHis)

Ub-GFP-TS

S-Ub^77N^

Ub

Complex (Ubp3^C469A^-Trs-TS + Bre5-Trs-10xHis + Ub^3G^)

Abbreviations:

Trs: Thrombin cleavage site

TS: Twin-strep-tag

ALFA: ALFA-tag

His: Poly-histidine-tag

Tev: Tev cleavage site

S: Strep-tag

**Sequences**

Ubp3 (ALFA-Ubp3-Trs-TS)

MPSRLEEELRRRLTEPMNMQDANKEESYSMYPKTSSPPPPTPTNMQIPIYQAPLQMYGYTQAPYLYPTQIPAYSFNMVNQNQPIYHQSGSPHHLPPQNNINGGSTTNNNNINKKKWHSNGITNNNGSSGNQGANSSGSGMSYNKSHTYHHNYSNNHIPMMASPNSGSNAGMKKQTNSSNGNGSSATSPSYSSYNSSSQYDLYKFDVTKLKNLKENSSNLIQLPLFINTTEAEFAAASVQRYELNMKALNLNSESLENSSVEKSSAHHHTKSHSIPKHNEEVKTETHGEEEDAHDKKPHASKDAHELKKKTEVKKEDAKQDRNEKVIQEPQATVLPVVDKKEPEESVEENTSKTSSPSPSPPAAKSWSAIASDAIKSRQASNKTVSGSMVTKTPISGTTAGVSSTNMAAATIGKSSSPLLSKQPQKKDKKYVPPSTKGIEPLGSIALRMCFDPDFISYVLRNKDVENKIPVHSIIPRGIINRANICFMSSVLQVLLYCKPFIDVINVLSTRNTNSRVGTSSCKLLDACLTMYKQFDKETYEKKFLENADDAEKTTESDAKKSSKSKSFQHCATADAVKPDEFYKTLSTIPKFKDLQWGHQEDAEEFLTHLLDQLHEELISAIDGLTDNEIQNMLQSINDEQLKVFFIRNLSRYGKAEFIKNASPRLKELIEKYGVINDDSTEENGWHEVSGSSKRGKKTKTAAKRTVEIVPSPISKLFGGQFRSVLDIPNNKESQSITLDPFQTIQLDISDAGVNDLETAFKKFSEYELLPFKSSSGNDVEAKKQTFIDKLPQVLLIQFKRFSFINNVNKDNAMTNYNAYNGRIEKIRKKIKYGHELIIPEESMSSITLKNNTSGIDDRRYKLTGVIYHHGVSSDGGHYTADVYHSEHNKWYRIDDVNITELEDDDVLKGGEEASDSRTAYILMYQKRNLVPRGSSGWSHPQFEKGGGSGGGSGGSSAWSHPQFEK

Ubp3^C469A^ (ALFA-Ubp3(C469A)-Trs-TS)

MPSRLEEELRRRLTEPMNMQDANKEESYSMYPKTSSPPPPTPTNMQIPIYQAPLQMYGYTQAPYLYPTQIPAYSFNMVNQNQPIYHQSGSPHHLPPQNNINGGSTTNNNNINKKKWHSNGITNNNGSSGNQGANSSGSGMSYNKSHTYHHNYSNNHIPMMASPNSGSNAGMKKQTNSSNGNGSSATSPSYSSYNSSSQYDLYKFDVTKLKNLKENSSNLIQLPLFINTTEAEFAAASVQRYELNMKALNLNSESLENSSVEKSSAHHHTKSHSIPKHNEEVKTETHGEEEDAHDKKPHASKDAHELKKKTEVKKEDAKQDRNEKVIQEPQATVLPVVDKKEPEESVEENTSKTSSPSPSPPAAKSWSAIASDAIKSRQASNKTVSGSMVTKTPISGTTAGVSSTNMAAATIGKSSSPLLSKQPQKKDKKYVPPSTKGIEPLGSIALRMCFDPDFISYVLRNKDVENKIPVHSIIPRGIINRANIAFMSSVLQVLLYCKPFIDVINVLSTRNTNSRVGTSSCKLLDACLTMYKQFDKETYEKKFLENADDAEKTTESDAKKSSKSKSFQHCATADAVKPDEFYKTLSTIPKFKDLQWGHQEDAEEFLTHLLDQLHEELISAIDGLTDNEIQNMLQSINDEQLKVFFIRNLSRYGKAEFIKNASPRLKELIEKYGVINDDSTEENGWHEVSGSSKRGKKTKTAAKRTVEIVPSPISKLFGGQFRSVLDIPNNKESQSITLDPFQTIQLDISDAGVNDLETAFKKFSEYELLPFKSSSGNDVEAKKQTFIDKLPQVLLIQFKRFSFINNVNKDNAMTNYNAYNGRIEKIRKKIKYGHELIIPEESMSSITLKNNTSGIDDRRYKLTGVIYHHGVSSDGGHYTADVYHSEHNKWYRIDDVNITELEDDDVLKGGEEASDSRTAYILMYQKRNLVPRGSSGWSHPQFEKGGGSGGGSGGSSAWSHPQFEK

Ubp3^Cat^ (6xHis-Tev-Ubp3(407-912)-Trs-TS)

MHHHHHHENLYFQGPQKKDKKYVPPSTKGIEPLGSIALRMCFDPDFISYVLRNKDVENKIPVHSIIPRGIINRANICFMSSVLQVLLYCKPFIDVINVLSTRNTNSRVGTSSCKLLDACLTMYKQFDKETYEKKFLENADDAEKTTESDAKKSSKSKSFQHCATADAVKPDEFYKTLSTIPKFKDLQWGHQEDAEEFLTHLLDQLHEELISAIDGLTDNEIQNMLQSINDEQLKVFFIRNLSRYGKAEFIKNASPRLKELIEKYGVINDDSTEENGWHEVSGSSKRGKKTKTAAKRTVEIVPSPISKLFGGQFRSVLDIPNNKESQSITLDPFQTIQLDISDAGVNDLETAFKKFSEYELLPFKSSSGNDVEAKKQTFIDKLPQVLLIQFKRFSFINNVNKDNAMTNYNAYNGRIEKIRKKIKYGHELIIPEESMSSITLKNNTSGIDDRRYKLTGVIYHHGVSSDGGHYTADVYHSEHNKWYRIDDVNITELEDDDVLKGGEEASDSRTAYILMYQKRNLVPRGSSGWSHPQFEKGGGSGGGSGGSSAWSHPQFEK

S-His-Tev-Ubp3(407-912)

MGSWSHPQFEKGSGMHHHHHHENLYFQGPQKKDKKYVPPSTKGIEPLGSIALRMCFDPDFISYVLRNKDVENKIPVHSIIPRGIINRANICFMSSVLQVLLYCKPFIDVINVLSTRNTNSRVGTSSCKLLDACLTMYKQFDKETYEKKFLENADDAEKTTESDAKKSSKSKSFQHCATADAVKPDEFYKTLSTIPKFKDLQWGHQEDAEEFLTHLLDQLHEELISAIDGLTDNEIQNMLQSINDEQLKVFFIRNLSRYGKAEFIKNASPRLKELIEKYGVINDDSTEENGWHEVSGSSKRGKKTKTAAKRTVEIVPSPISKLFGGQFRSVLDIPNNKESQSITLDPFQTIQLDISDAGVNDLETAFKKFSEYELLPFKSSSGNDVEAKKQTFIDKLPQVLLIQFKRFSFINNVNKDNAMTNYNAYNGRIEKIRKKIKYGHELIIPEESMSSITLKNNTSGIDDRRYKLTGVIYHHGVSSDGGHYTADVYHSEHNKWYRIDDVNITELEDDDVLKGGEEASDSRTAYILMYQKRN

Ubp3^Cat^ (Tags removed for experiments)

GPQKKDKKYVPPSTKGIEPLGSIALRMCFDPDFISYVLRNKDVENKIPVHSIIPRGIINRANICFMSSVLQVLLYCKPFIDVINVLSTRNTNSRVGTSSCKLLDACLTMYKQFDKETYEKKFLENADDAEKTTESDAKKSSKSKSFQHCATADAVKPDEFYKTLSTIPKFKDLQWGHQEDAEEFLTHLLDQLHEELISAIDGLTDNEIQNMLQSINDEQLKVFFIRNLSRYGKAEFIKNASPRLKELIEKYGVINDDSTEENGWHEVSGSSKRGKKTKTAAKRTVEIVPSPISKLFGGQFRSVLDIPNNKESQSITLDPFQTIQLDISDAGVNDLETAFKKFSEYELLPFKSSSGNDVEAKKQTFIDKLPQVLLIQFKRFSFINNVNKDNAMTNYNAYNGRIEKIRKKIKYGHELIIPEESMSSITLKNNTSGIDDRRYKLTGVIYHHGVSSDGGHYTADVYHSEHNKWYRIDDVNITELEDDDVLKGGEEASDSRTAYILMYQKRN

Bre5 (Bre5-Trs-10xHis)

MGVTVQDICFAFLQNYYERMRTDPSKLAYFYASTAELTHTNYQSKSTNEKDDVLPTVKVTGRENINKFFSRNDAKVRSLKLKLDTIDFQYTGHLHKSILIMATGEMFWTGTPVYKFCQTFILLPSSNGSTFDITNDIIRFISNSFKPYVLTDASLSQSNEENSVSAVEEDKIRHESGVEKEKEKEKSPEISKPKAKKETVKDTTAPTESSTQEKPIVDHSQPRAIPVTKESKIHTETVPSSTKGNHKQDEVSTEELGNVTKLNEKSHKAEKKAAPIKTKEGSVEAINAVNNSSLPNGKEVSDEKPVPGGVKEAETEIKPIEPQVSDAKESGNNASTPSSSPEPVANPPKMTWASKLMNENSDRISKNNTTVEYIRPETLPKKPTERKFEMGNRRDNASANSKNKKKPVFSTVNKDGFYPIYIRGTNGLREEKLRSALEKEFGKVMRITAADNFAVVDFETQKSQIDALEKKKKSIDGIEVCLERKTVKKPTSNNPPGIFTNGTRSHRKQPLKRKDLVPRGSSSGHHHHHHHHHH

Ub-GFP-TS

MQIFVKTLTGKTITLEVESSDTIDNVKSKIQDKEGIPPDQQRLIFAGKQLEDGRTLSDYNIQKESTLHLVLRLRGGVSKGEELFTGVVPILVELDGDVNGHKFSVSGEGEGDATYGKLTLKFICTTGKLPVPWPTLVTTLTYGVQCFSRYPDHMKQHDFFKSAMPEGYVQERTIFFKDDGNYKTRAEVKFEGDTLVNRIELKGIDFKEDGNILGHKLEYNYNSHNVYIMADKQKNGIKVNFKIRHNIEDGSVQLADHYQQNTPIGDGPVLLPDNHYLSTQSALSKDPNEKRDHMVLLEFVTAAGITLGMDELYKLVPRGSSGWSHPQFEKGGGSGGGSGGSSAWSHPQFEK

S-Ub^77N^

MGSWSHPQFEKQIFVKTLTGKTITLEVESSDTIDNVKSKIQDKEGIPPDQQRLIFAGKQLEDGRTLSDYNIQKESTLHLVLRLRGGN

Ub

MQIFVKTLTGKTITLEVESSDTIDNVKSKIQDKEGIPPDQQRLIFAGKQLEDGRTLSDYNIQKESTLHLVLRLRGG

Complex (Ubp3^C469A^-Trs-TS + Bre5-Trs-10xHis + Ub3G)

MNMQDANKEESYSMYPKTSSPPPPTPTNMQIPIYQAPLQMYGYTQAPYLYPTQIPAYSFNMVNQNQPIYHQSGSPHHLPPQNNINGGSTTNNNNINKKKWHSNGITNNNGSSGNQGANSSGSGMSYNKSHTYHHNYSNNHIPMMASPNSGSNAGMKKQTNSSNGNGSSATSPSYSSYNSSSQYDLYKFDVTKLKNLKENSSNLIQLPLFINTTEAEFAAASVQRYELNMKALNLNSESLENSSVEKSSAHHHTKSHSIPKHNEEVKTETHGEEEDAHDKKPHASKDAHELKKKTEVKKEDAKQDRNEKVIQEPQATVLPVVDKKEPEESVEENTSKTSSPSPSPPAAKSWSAIASDAIKSRQASNKTVSGSMVTKTPISGTTAGVSSTNMAAATIGKSSSPLLSKQPQKKDKKYVPPSTKGIEPLGSIALRMCFDPDFISYVLRNKDVENKIPVHSIIPRGIINRANIAFMSSVLQVLLYCKPFIDVINVLSTRNTNSRVGTSSCKLLDACLTMYKQFDKETYEKKFLENADDAEKTTESDAKKSSKSKSFQHCATADAVKPDEFYKTLSTIPKFKDLQWGHQEDAEEFLTHLLDQLHEELISAIDGLTDNEIQNMLQSINDEQLKVFFIRNLSRYGKAEFIKNASPRLKELIEKYGVINDDSTEENGWHEVSGSSKRGKKTKTAAKRTVEIVPSPISKLFGGQFRSVLDIPNNKESQSITLDPFQTIQLDISDAGVNDLETAFKKFSEYELLPFKSSSGNDVEAKKQTFIDKLPQVLLIQFKRFSFINNVNKDNAMTNYNAYNGRIEKIRKKIKYGHELIIPEESMSSITLKNNTSGIDDRRYKLTGVIYHHGVSSDGGHYTADVYHSEHNKWYRIDDVNITELEDDDVLKGGEEASDSRTAYILMYQKRNLVPRGSSGWSHPQFEKGGGSGGGSGGSSAWSHPQFEKDPVFAC

MGVTVQDICFAFLQNYYERMRTDPSKLAYFYASTAELTHTNYQSKSTNEKDDVLPTVKVTGRENINKFFSRNDAKVRSLKLKLDTIDFQYTGHLHKSILIMATGEMFWTGTPVYKFCQTFILLPSSNGSTFDITNDIIRFISNSFKPYVLTDASLSQSNEENSVSAVEEDKIRHESGVEKEKEKEKSPEISKPKAKKETVKDTTAPTESSTQEKPIVDHSQPRAIPVTKESKIHTETVPSSTKGNHKQDEVSTEELGNVTKLNEKSHKAEKKAAPIKTKEGSVEAINAVNNSSLPNGKEVSDEKPVPGGVKEAETEIKPIEPQVSDAKESGNNASTPSSSPEPVANPPKMTWASKLMNENSDRISKNNTTVEYIRPETLPKKPTERKFEMGNRRDNASANSKNKKKPVFSTVNKDGFYPIYIRGTNGLREEKLRSALEKEFGKVMRITAADNFAVVDFETQKSQIDALEKKKKSIDGIEVCLERKTVKKPTSNNPPGIFTNGTRSHRKQPLKRKDLVPRGSSSGHHHHHHHHHH

MQIFVKTLTGKTITLEVESSDTIDNVKSKIQDKEGIPPDQQRLIFAGKQLEDGRTLSDYNIQKESTLHLVLRLRGGGGG

**Table S1: Constructs, their molecular weight (in Da) and extinction coefficient as calculated from amino acid sequence in Benchling.**

| Protein | M_W_ (Da) | 𝞮 (M^-1^cm^-1^) (Cysteines oxidized) |
| --- | --- | --- |
| ALFA-Ubp3-Trs-TS | 107,620.21 | 92,515.00 |
| ALFA-Ubp3(C469A)-Trs-TS | 107,588.15 | 92,390.00 |
| 6xHis-Tev-Ubp3(407-912)-Trs-TS | 63,231.68 | 57,675.00 |
| S-His-Tev-Ubp3(407-912)  Ubp3(407-912) (no tags) | 61,038.37 | 52,175.00 |
|  | 57,772.79 | 45,185.00 |
| Bre5-Trs-10xHis | 59,885.90 | 27,515.00 |
| Ub-GFP-TS | 39,058.83 | 34,505.00 |
| S-Ub^77N^ | 9,855.10 | 6,990.00 |
| Ub | 8,556.72 | 1,490.00 |
| Ubp3^C469A^-Trs-TS + Bre5-Trs-10xHis + Ub^3G^ | 174,840.38 (1:1:1 stoichiometry) | Evaluated with 1 A_280_ = 1 mg/ml |

**Strains**

∆malP *E. coli* (Keio Collection)

∆elaD *E. coli* (Keio Collection)

DH5ɑ *E. coli*

BL21(DE3) *E. coli*

**Table S2: Overview of the different constructs, expression strains and duration, purification columns and concentrators used during purification.**

| Construct | Strain and expression time | 1^st^ chromatography | 2^nd^ chromatography | Concentrator (Mw cutoff kDa) | SEC column |
| --- | --- | --- | --- | --- | --- |
| ALFA-Ubp3-TS | ∆malP  3 h | StrepTrap XT 5 ml | ALFA Selector CE approximately 800 µl resin | 50 | Superose 6 Increase 10/300 GL |
| ALFA-Ubp3^C469A^-TS | ∆malP  3 h | StrepTrap XT 5 ml | ALFA Selector CE approximately 800 µl resin | 50 | Superose 6 Increase 10/300 GL |
| His-Ubp3^Cat^-TS | ∆malP  3 h | HiTrap™ IMAC HP 5 ml | StrepTrap XT 5 ml | 30 | HiLoad 16/600 Superdex 200 pg |
| SH-Ubp3^Cat^ | ∆malP  overnight | HiTrap™ IMAC HP 5 ml | StrepTrap XT 5 ml | 30 | Superdex 200 Increase 10/300 GL |
| Bre5-His | ∆malP  overnight | HiTrap™ Chelating HP 5 ml | HiTrap™ SP HP 5 ml | 30 | Superose 6 Increase 10/300 GL |
| Ub-GFP-TS | Bl21(DE3)  2 h | StrepTrap XT 5 ml | - | 30 | Superdex 200 Increase 10/300 GL |
| S-Ub^77N^ | ∆elaD  overnight | StrepTrap XT 5 ml | - | 3 | HiLoad 16/600 Superdex 75 pg |
| Ub | Bl21(DE3)  overnight | HiTrap SP FF 5 ml | - | 3 | HiLoad 16/600 Superdex 75 pg |
| Full complex | ∆malP  3 h | HiTrap™ IMAC HP 5 ml | StrepTrap XT 5 ml | 50 | Superose 6 Increase 10/300 GL |

**Table S3: Proteins and the corresponding primary and secondary antibodies used for detection.**

| Construct | 1^st^ Antibody | 2^nd^ Antibody |
| --- | --- | --- |
| ALFA-Ubp3-TS / ALFA-Ubp3^C469A^-TS | Affinity purified rabbit anti-ALFA polyclonal antibody (NanoTag Biotechnologies) | Anti-Rabbit IgG (whole molecule)–Peroxidase conjugated (Sigma) |
|  | Strep Tag II Monoclonal Antibody (Millipore) | Peroxidase-AffiniPure Goat Anti-mouse IgG (H+L) (Jackson ImmunoResearch Laboratories) |
| His-Ubp3^Cat^-TS / SH-Ubp3^Cat^ | Strep Tag II Monoclonal Antibody (Millipore) | Peroxidase-AffiniPure Goat Anti-mouse IgG (H+L) (Jackson ImmunoResearch Laboratories) |
|  | Penta-His Antibody, BSA-free (Qiagen) | Peroxidase-AffiniPure Goat Anti-mouse IgG (H+L) (Jackson ImmunoResearch Laboratories) |
| Bre5-His | Penta-His Antibody, BSA-free (Qiagen) | Peroxidase-AffiniPure Goat Anti-mouse IgG (H+L) (Jackson ImmunoResearch Laboratories) |
| Ub-GFP-TS | Strep Tag II Monoclonal Antibody (Millipore) | Peroxidase-AffiniPure Goat Anti-mouse IgG (H+L) (Jackson ImmunoResearch Laboratories) |
| Ub / S-Ub^77N^ | Anti-Ubiquitin antibody (ab19247) (abcam) | Anti-Rabbit IgG (whole molecule)–Peroxidase conjugated (Sigma) |
| Full complex | Strep Tag II Monoclonal Antibody (Millipore) | Peroxidase-AffiniPure Goat Anti-mouse IgG (H+L) (Jackson ImmunoResearch Laboratories) |
|  | Penta-His Antibody, BSA-free (Qiagen) | Peroxidase-AffiniPure Goat Anti-mouse IgG (H+L) (Jackson ImmunoResearch Laboratories) |
|  | Anti-Ubiquitin antibody (ab19247) (abcam) | Anti-Rabbit IgG (whole molecule)–Peroxidase conjugated (Sigma) |

**Figures**

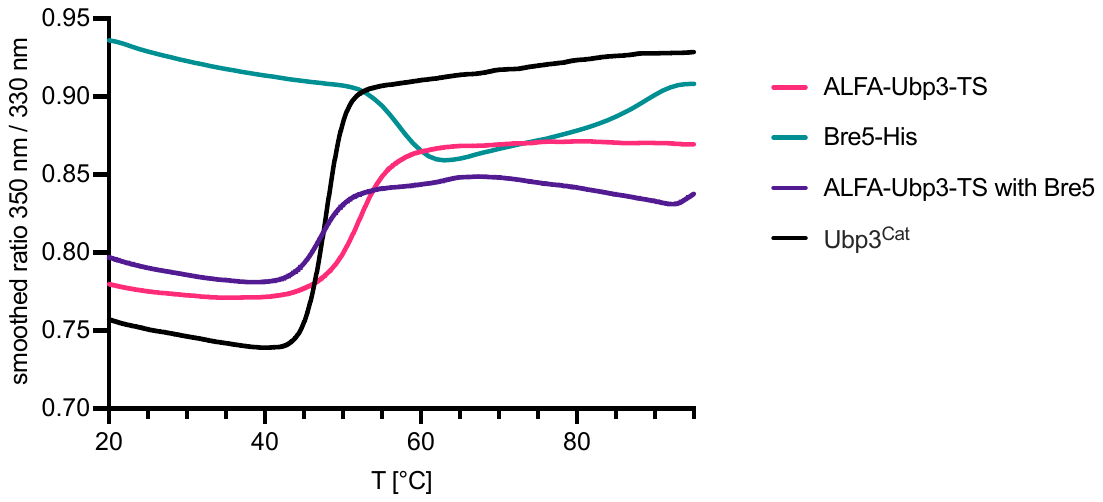

Figure S1: NanoDSF unfolding ratios measured for ALFA-Ubp3-TS (red), Bre5-His (cyan), both proteins mixed together prior to measurement (purple) and Ubp3^Cat^ (black). N = 2

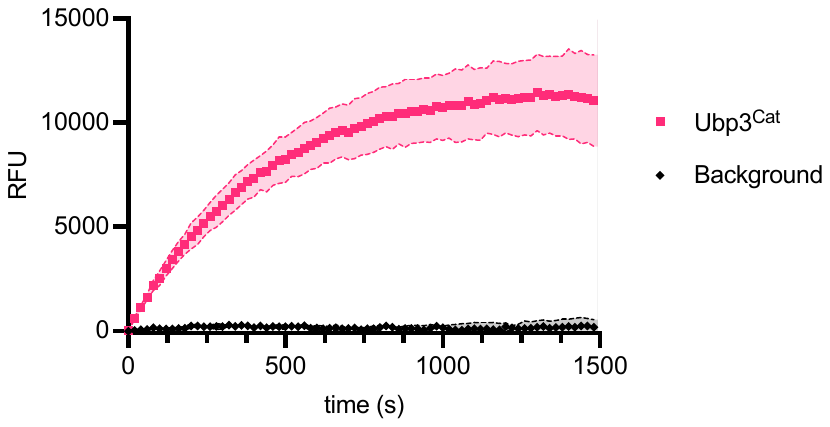

Figure S2: Ub-AMC cleavage assay, fluorescence increase over time in RFU for His-Ubp3^Cat^-TS (red squares) and background (black diamonds), shaded areas show the SD of three replicates. (Reaction was carried out as described in method section with 40 nM enzyme)

### SEC-SAXS results: BRE5-His

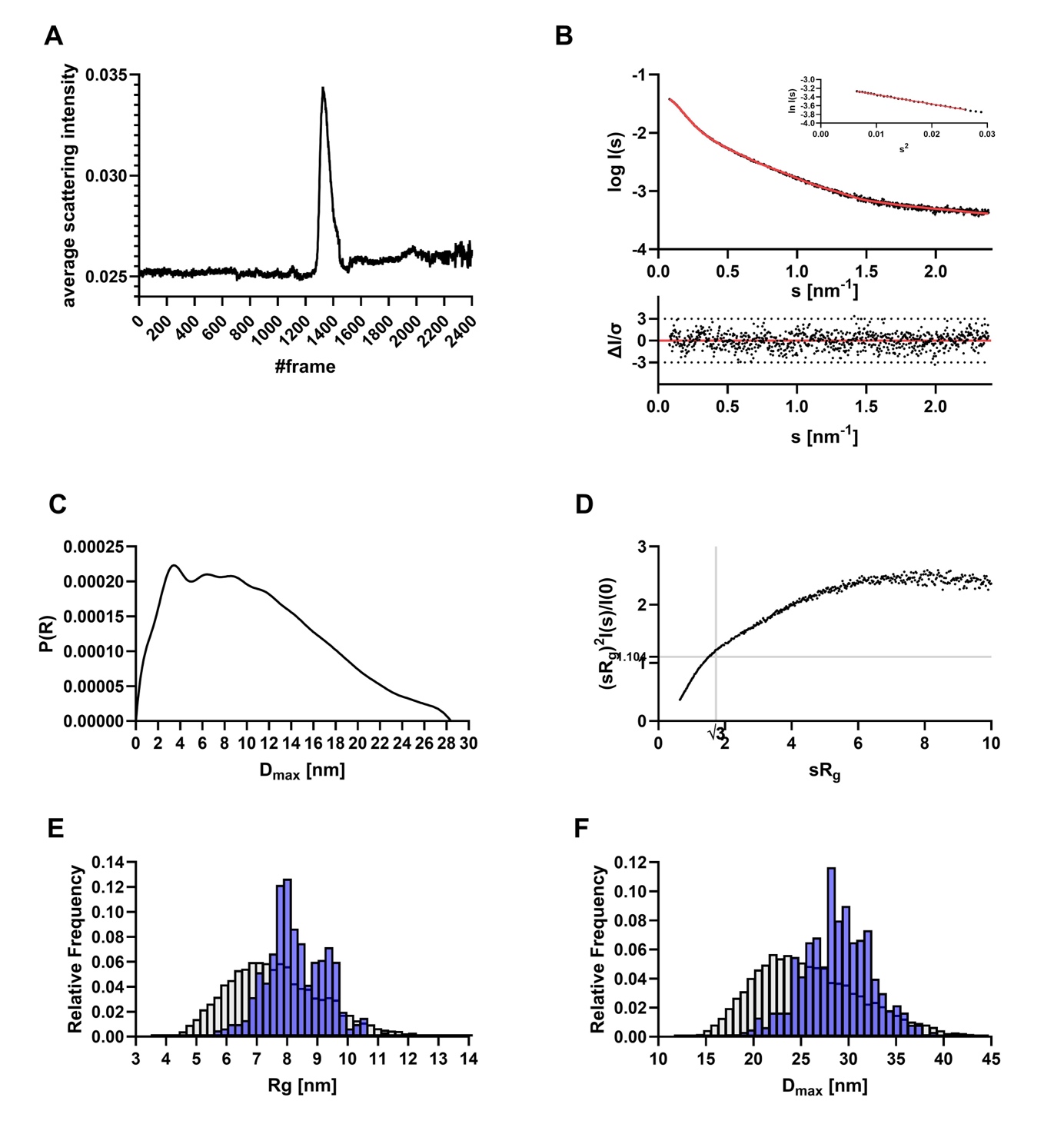

**Figure S3: Small-angle X-ray scattering data of Bre5-His. A:** CHROMIXS SEC SAXS elution profiles of Bre5. **B:** Experimental data are shown in black dots, with grey error bars. The EOM ensemble model fit (χ 2 value of 1.098) is shown as red line and below is the residual plot of the data The Guinier plot is added in the right corner. **C:** The *p(r)* function of Bre5 showed an elongated particle. **D:** The Dimensionless Kratky plot of showed an elongated particle with a high degree of flexibility. **F & G:** *R_g_* and *D_max_* distribution of Bre5. Ensemble random pool is shown in grey, selected EOM models are shown in blue.

### BRE5-His EOM results

\# Filename	Rg [nm]	Dmax [nm]	Fraction

1) ranch01505	8.18	30.45	~0.33 ( 2/ 6)

2) ranch02733	9.90	35.69	~0.17 ( 1/ 6)

3) ranch04859	8.04	28.45	~0.17 ( 1/ 6)

4) ranch08708	8.26	28.77	~0.33 ( 2/ 6)

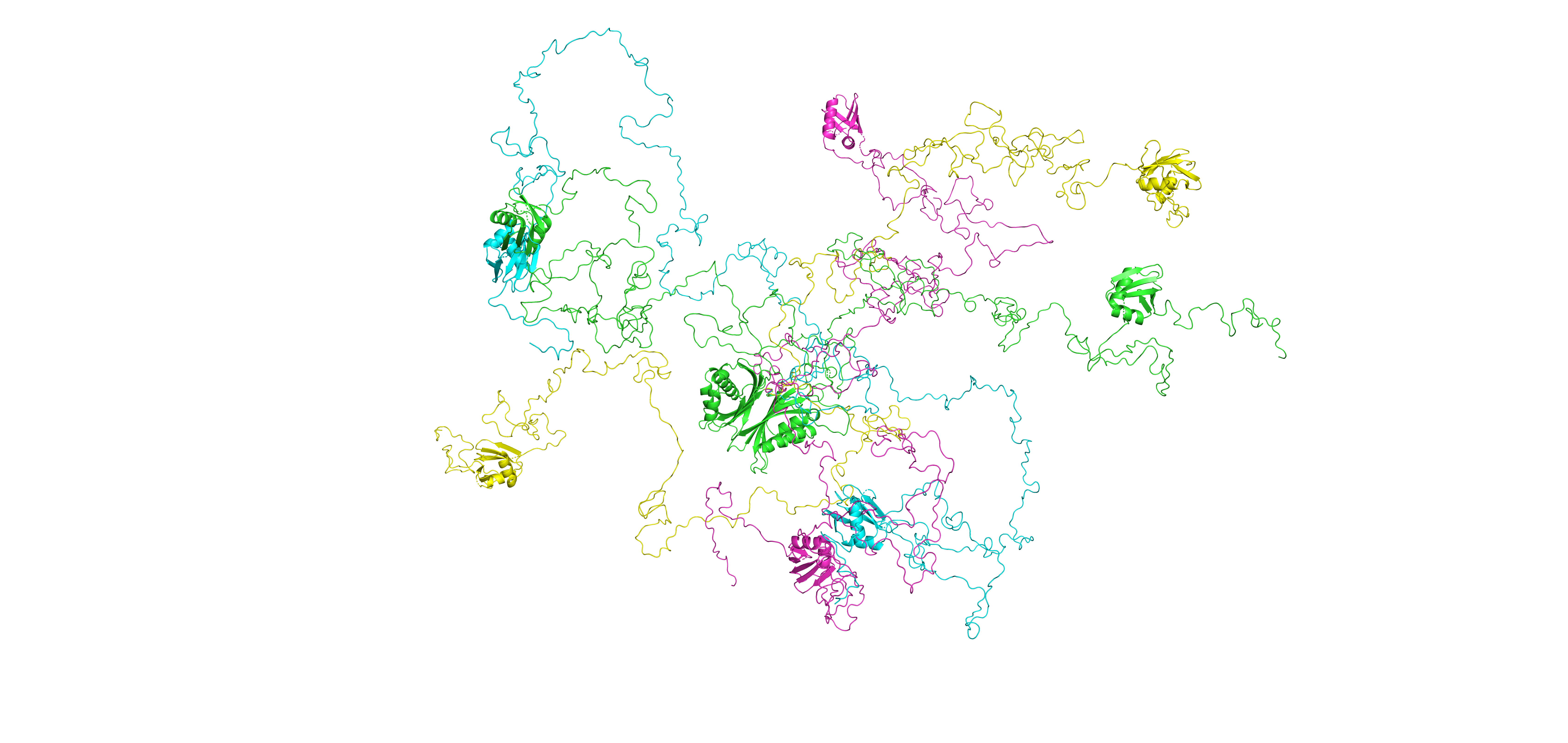

### Figure S4: Overlay of the selected EOM models Bre5. The ranch01505 model is shown in green. The ranch02733 model is shown in yellow. The ranch04859 model is shown in cyan and the ranch08708 model is shown in magenta.

### SEC-SAXS results: ALFA-Ubp3-Trs-TS

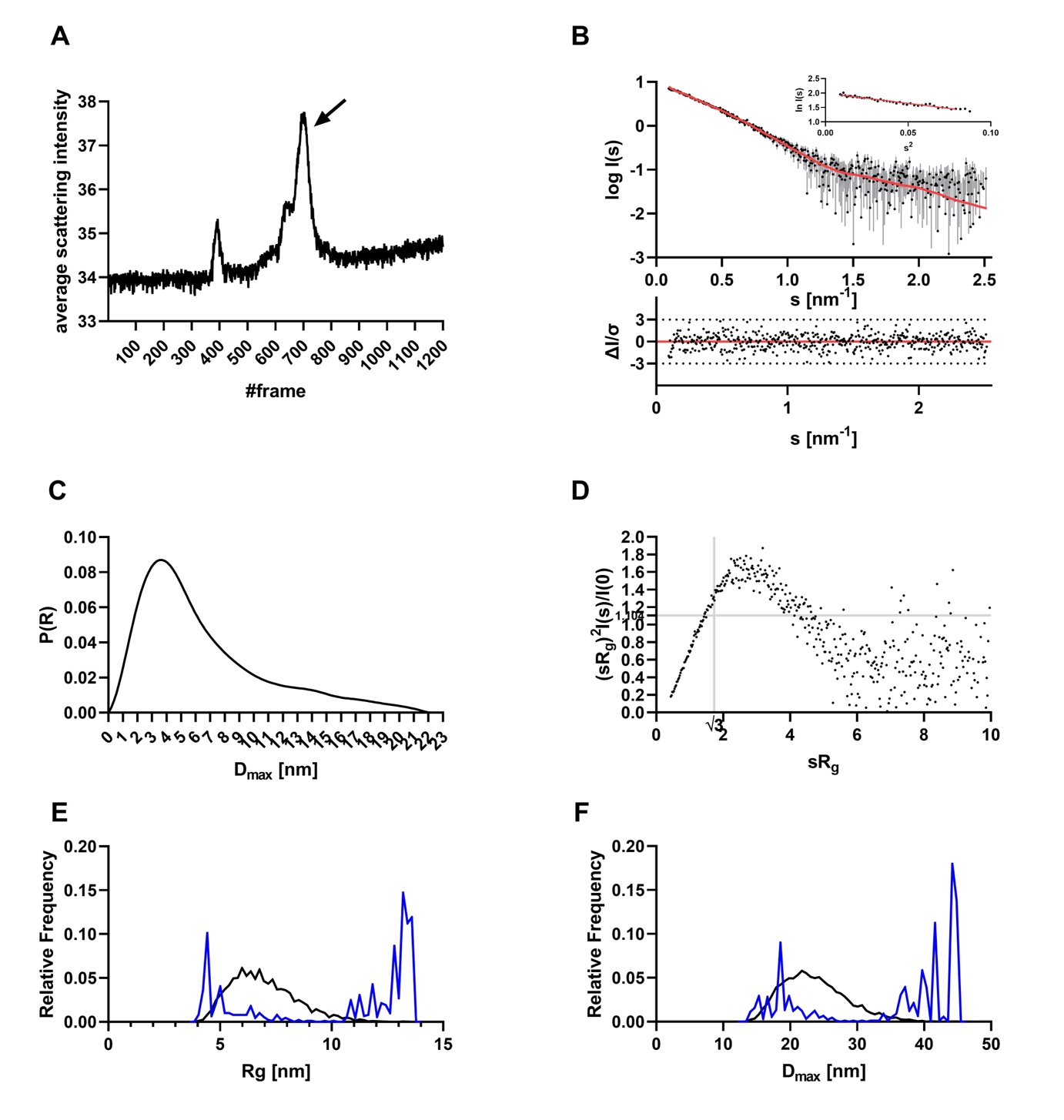

**Figure S5: Small-angle X-ray scattering data from ALFA-Ubp3-TS apo. A:** Chromixs SEC SAXS elution profile of ALFA-Ubp3-Trs-TS. **B:** Experimental data are shown in black dots, with grey error bars. The EOM fit (χ 2 value of 0.959) is shown as red line and below is the residual plot of the data. The Guinier plot is added in the right corner. **C:** The Distance distribution function *(p(r)* function) indicate an elongated molecule. **D:** Dimensionless Kratky plot indicate an elongated molecule with certain amounts of flexibility. **E & F:** *R_g_* and *D_max_* distribution of ALFA-Ubp3-Trs-TS. Ensemble random pool is shown in grey, selected EOM models are shown in blue.

**ALFA-Ubp3-Trs-TS EOM results**

\# Filename	Rg [nm]	Dmax [nm]	Fraction

1) ranch00190	13.31	44.87	~0.22 ( 2/ 9)

2) ranch03221	13.32	44.79	~0.44 ( 4/ 9)

3) ranch07712	4.59	18.71	~0.33 ( 3/ 9)

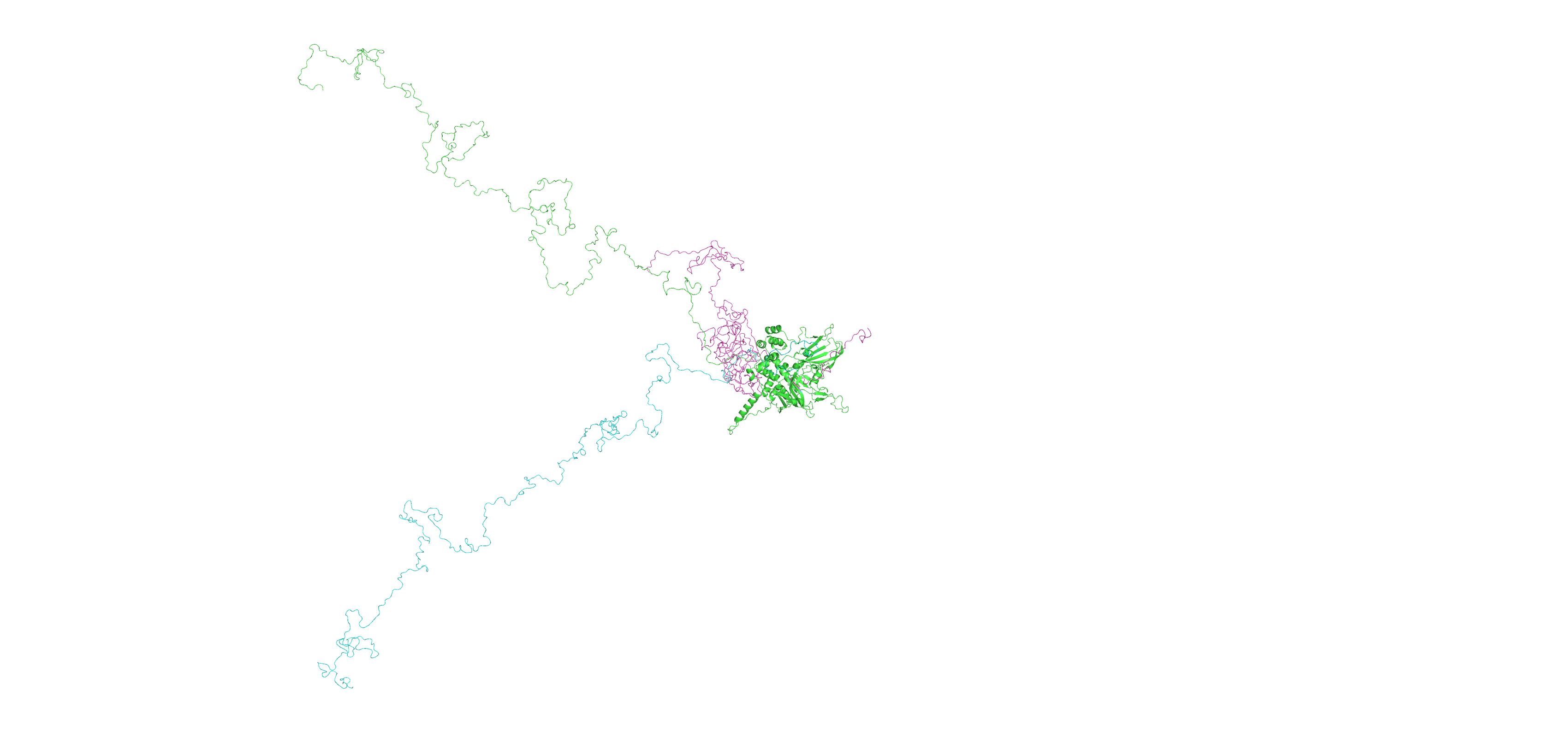

**Fig S6: ALFA-Ubp3-Trs-TS EOM models.** The AlphaFold3 model core is shown in green cartoon. The N-terminal tail was remodeled with EOM. Model ranch00190 is shown in green, ranch03221 in cyan and ranch07712 is magenta.

### SEC-SAXS results: 6xHis-Tev-Ubp3(407-912)-Trs-TS

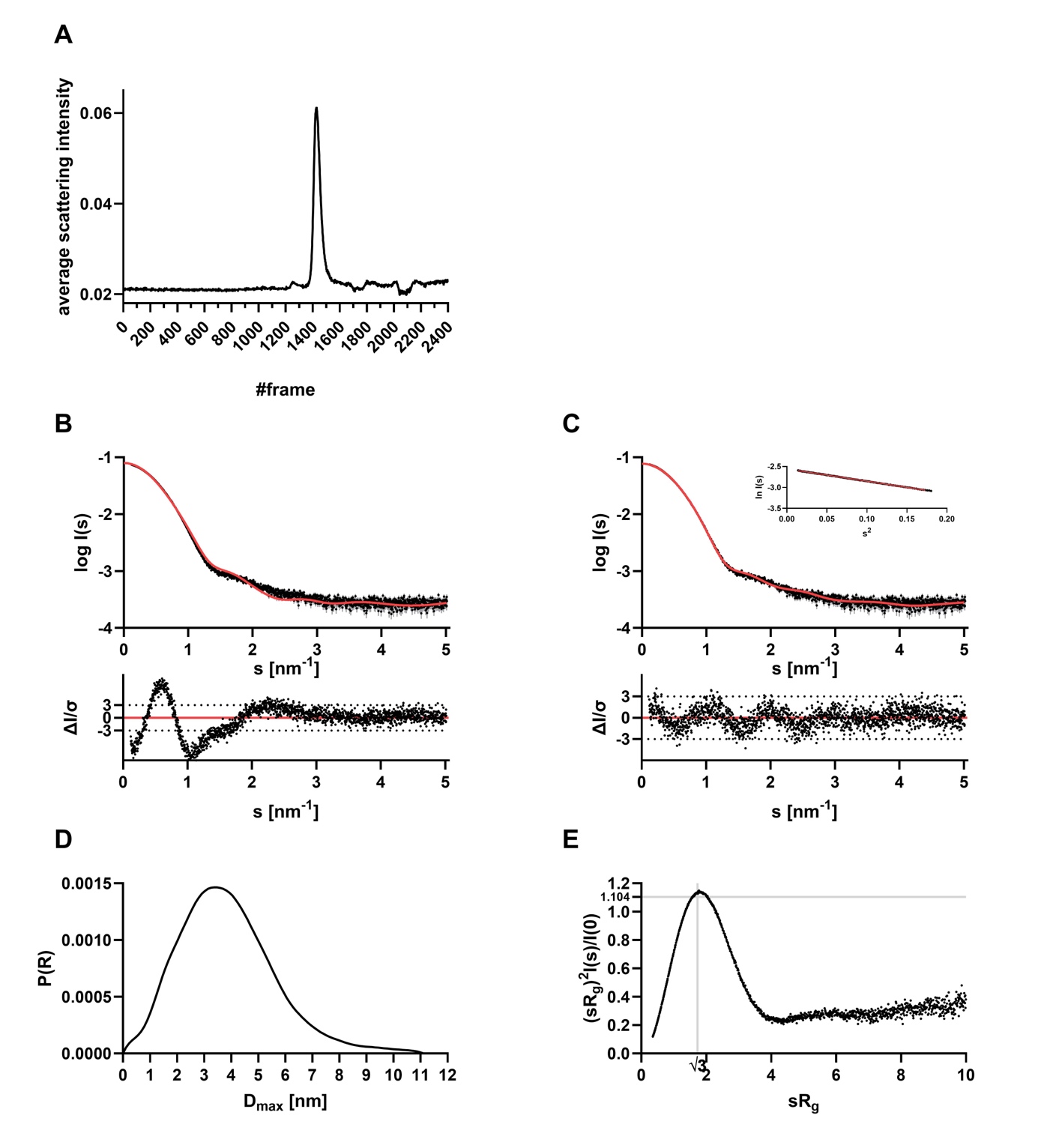

**Figure S7: Small-angle X-ray scattering data of Ubp3 catalytic domain. A:** CHROMIXS SEC SAXS elution profile of Ubp3 catalytic domain. **B:** Experimental data are shown in black dots, with grey error bars. The CRYSOL comparison fit (χ 2 value of 11.53) with the initial AF3 model is shown as red line and below is the residual plot of the data. **C:** Experimental data are shown in black dots, with grey error bars. The CRYSOL comparison fit with the best fit AFflecto model (χ 2 value of 1.52) is shown as red line and below is the residual plot of the data. The Guinier plot is added in the right corner. **D:** The Distance distribution function (*p(r)* function) indicate an elongated molecule. **E:** Dimensionless Kratky plot indicate a globular molecule with certain amounts of flexibility.

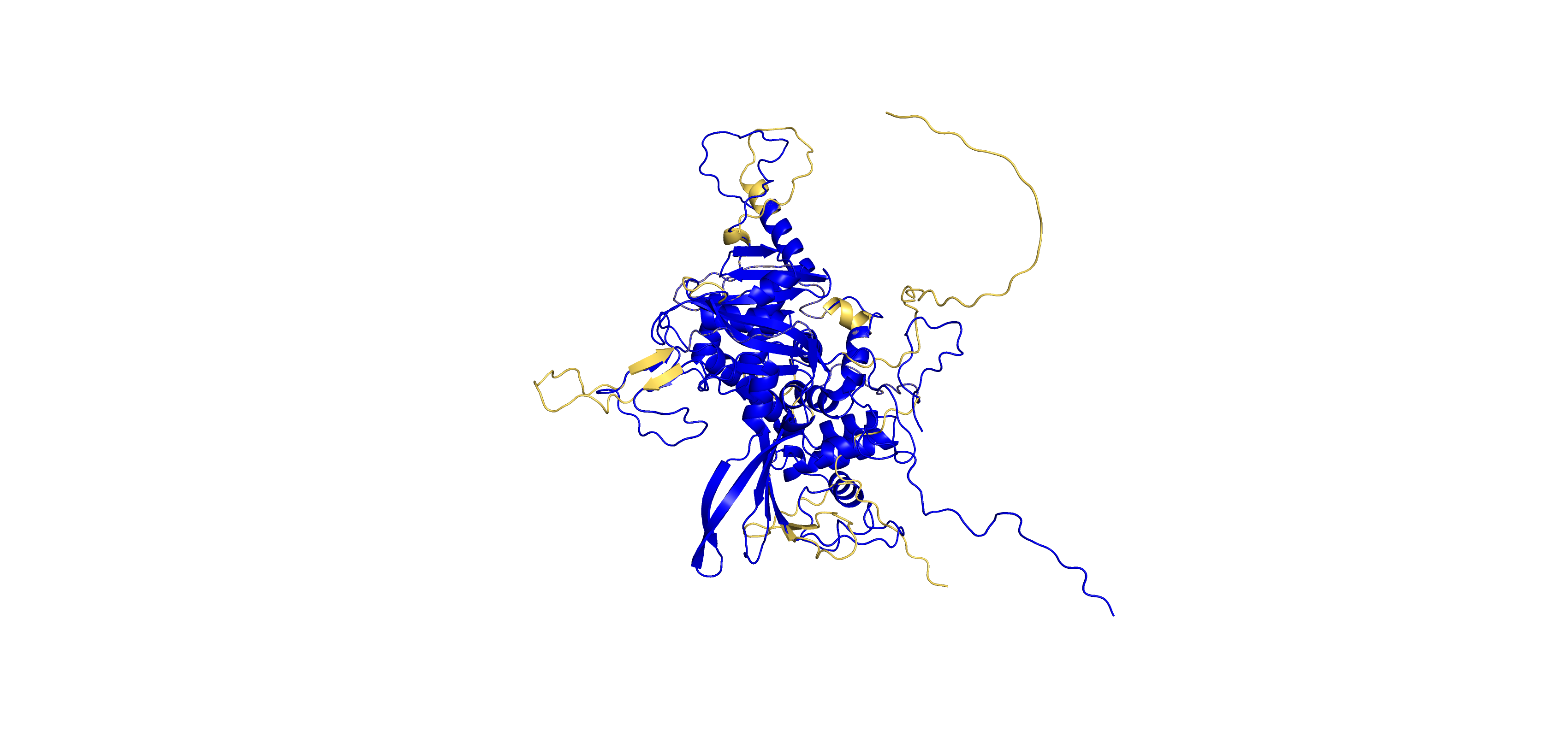

**Fig S8: Ubp3 catalytic domain:** Shown in gold is the original AF3 model and the best fit AFflecto model in blue. Flexible loop regions with low pIDDT score were sampled with different conformations.

### SEC-SAXS results: Bre5-Ubp3-Ubiquitin complex

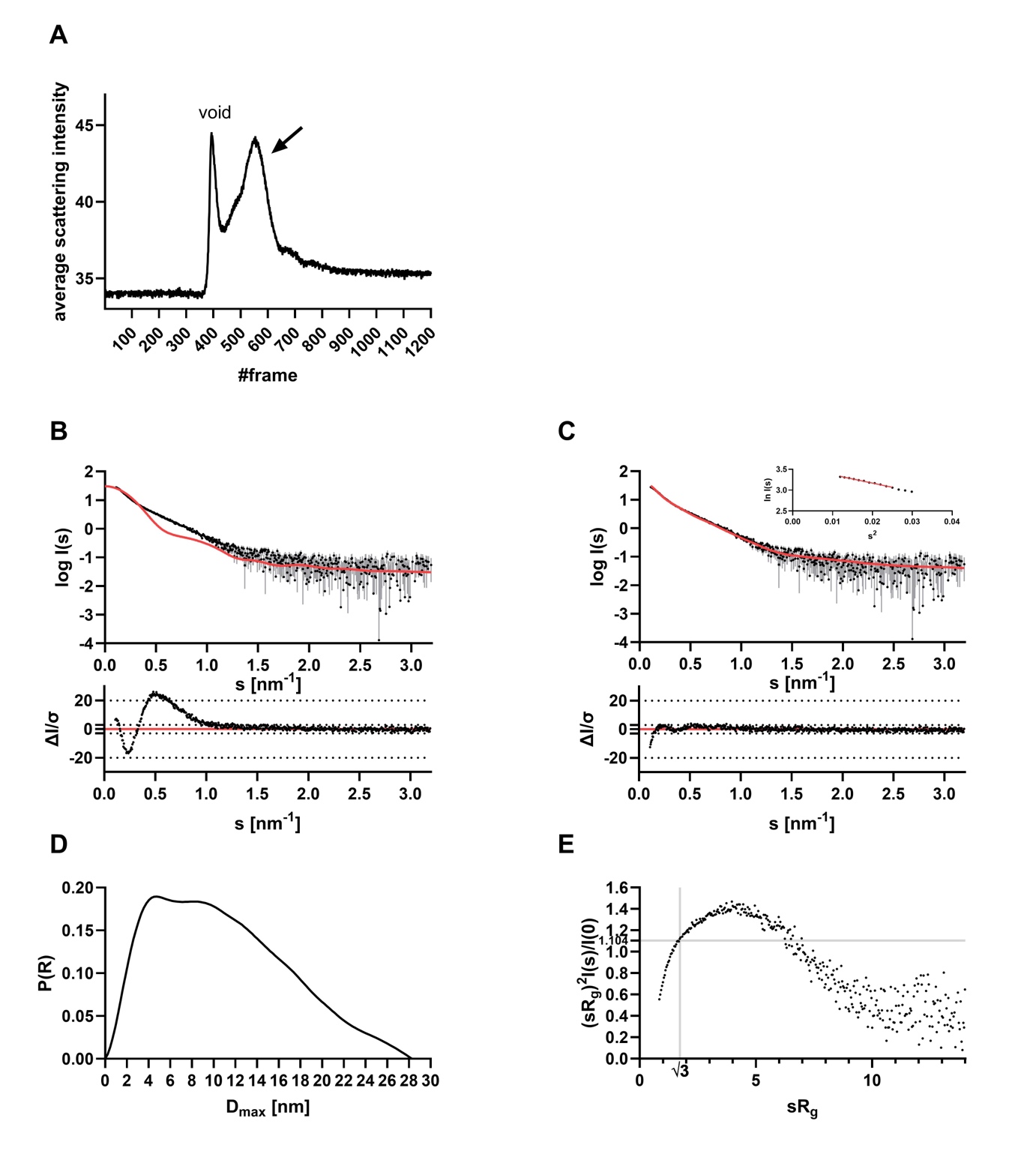

**Figure S9: Small-angle X-ray scattering data from the Bre5-Ubp3-Ubiquitin complex. A:** Chromixs SEC SAXS elution profile of the Bre5-Ubp3-Ubiquitin complex. **B:** Experimental data are shown in black dots, with grey error bars. The theoretical scattering fit of the initial AlphaFold3 model, created with CRYSOL (χ 2 value of 64.86), is shown as red line and below is the residual plot of the data. **C:** Experimental data are shown in black dots, with grey error bars. The BilboMD 2-state model fit (χ 2 value of 2.48) is shown as red line and below is the residual plot of the data. The Guinier plot is added in the right corner. **D:** The Distance distribution function *(p(r)* function) indicate an elongated multidomain molecule. **E:** Dimensionless Kratky plot indicate an elongated molecule with certain amounts of flexibility.

**BilboMD 3 state model result**

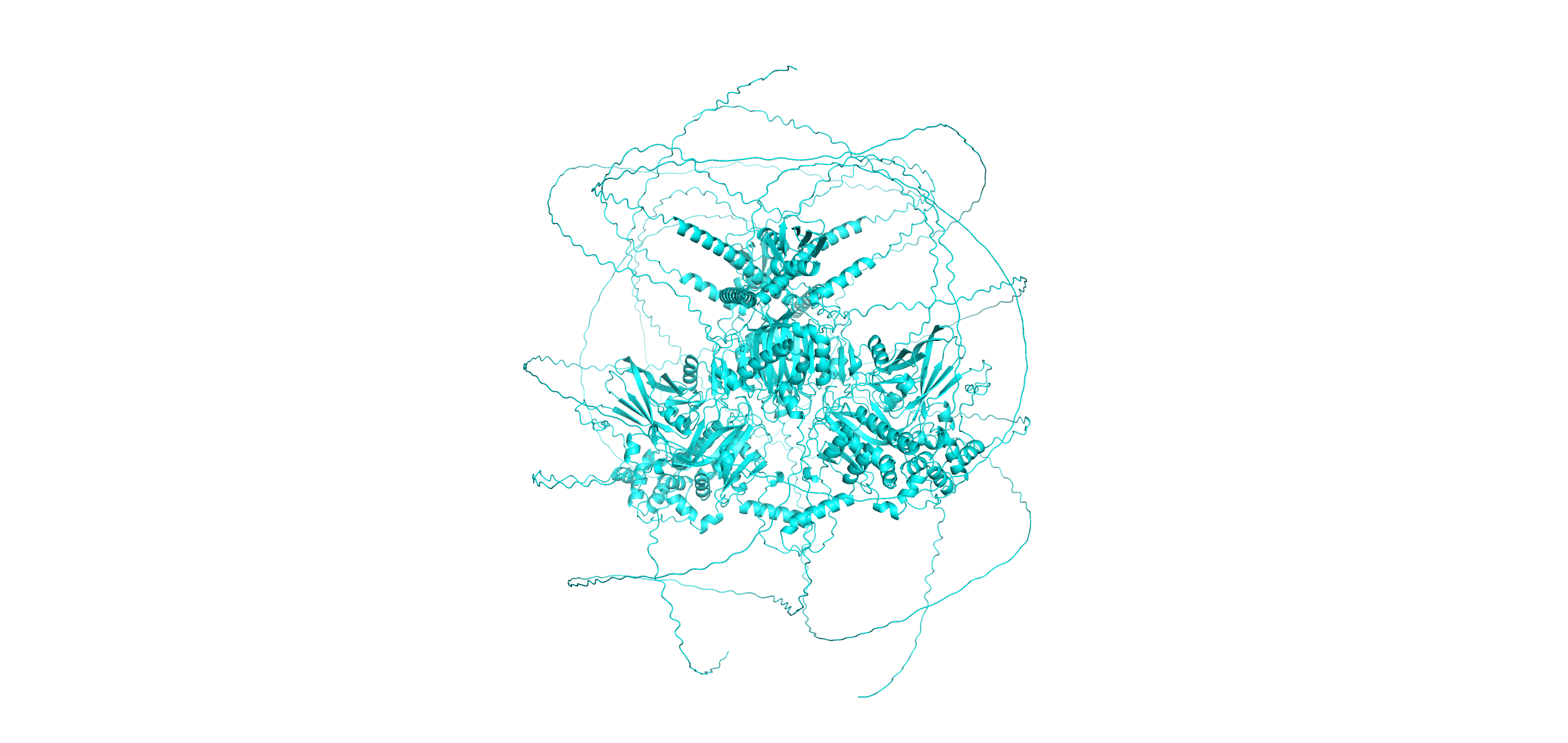

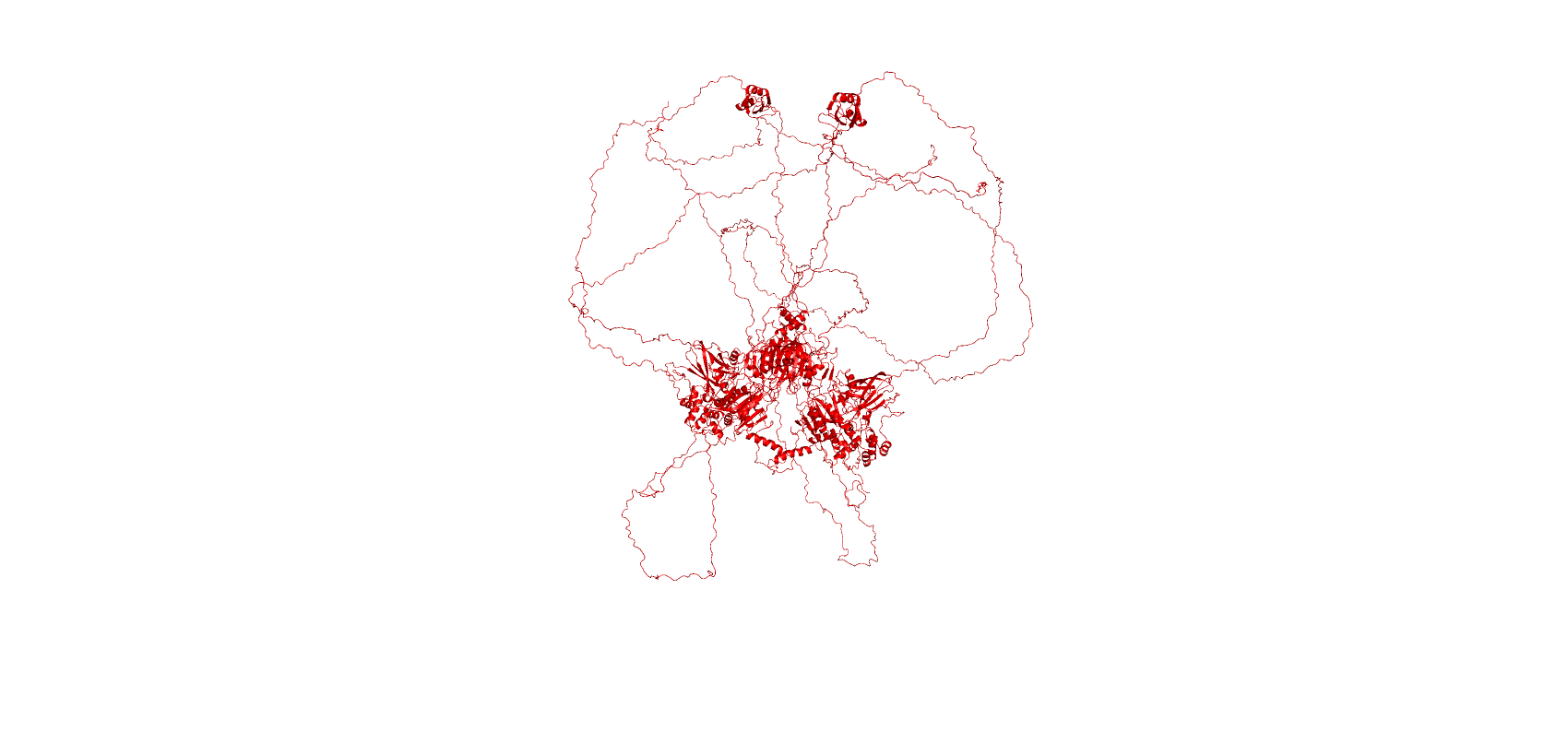

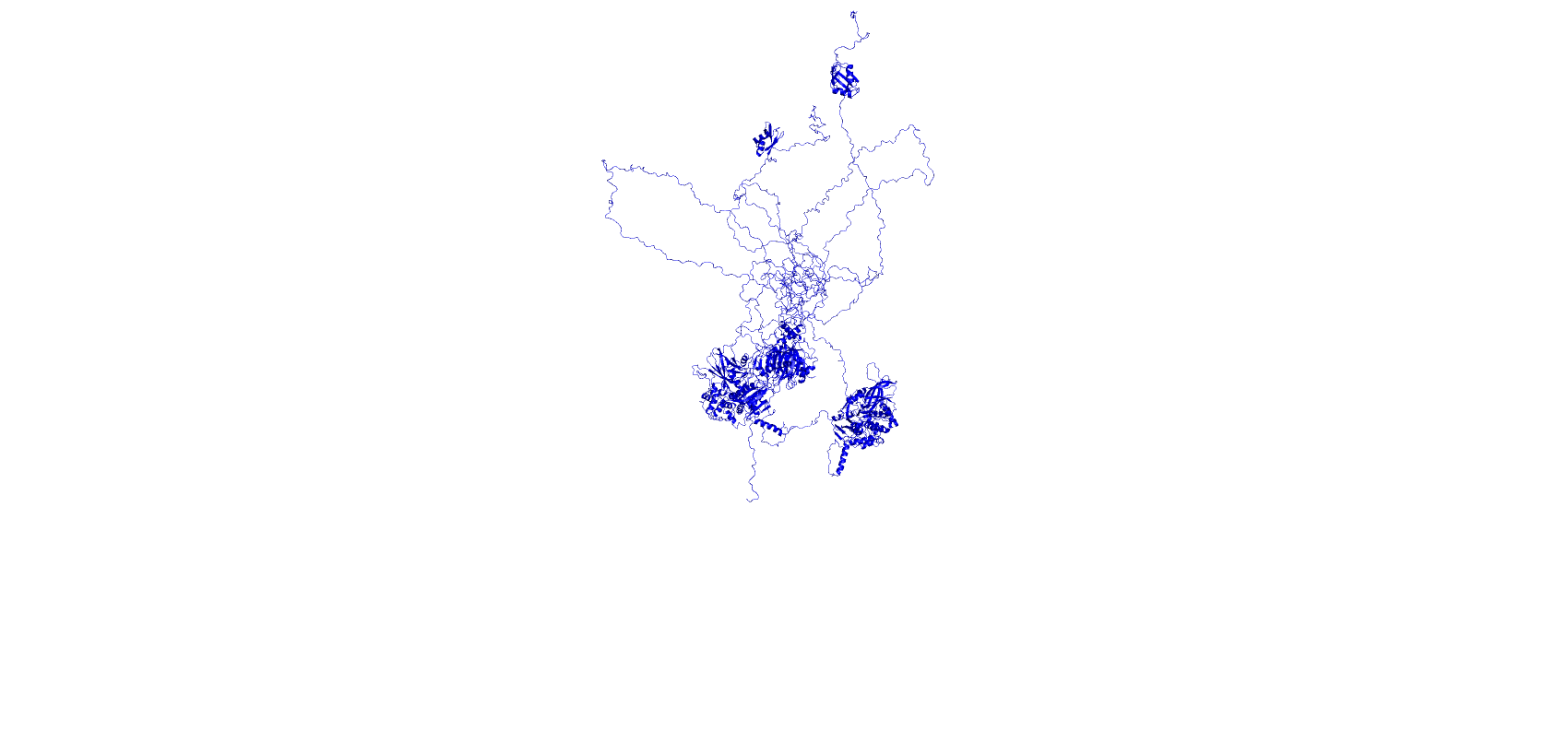

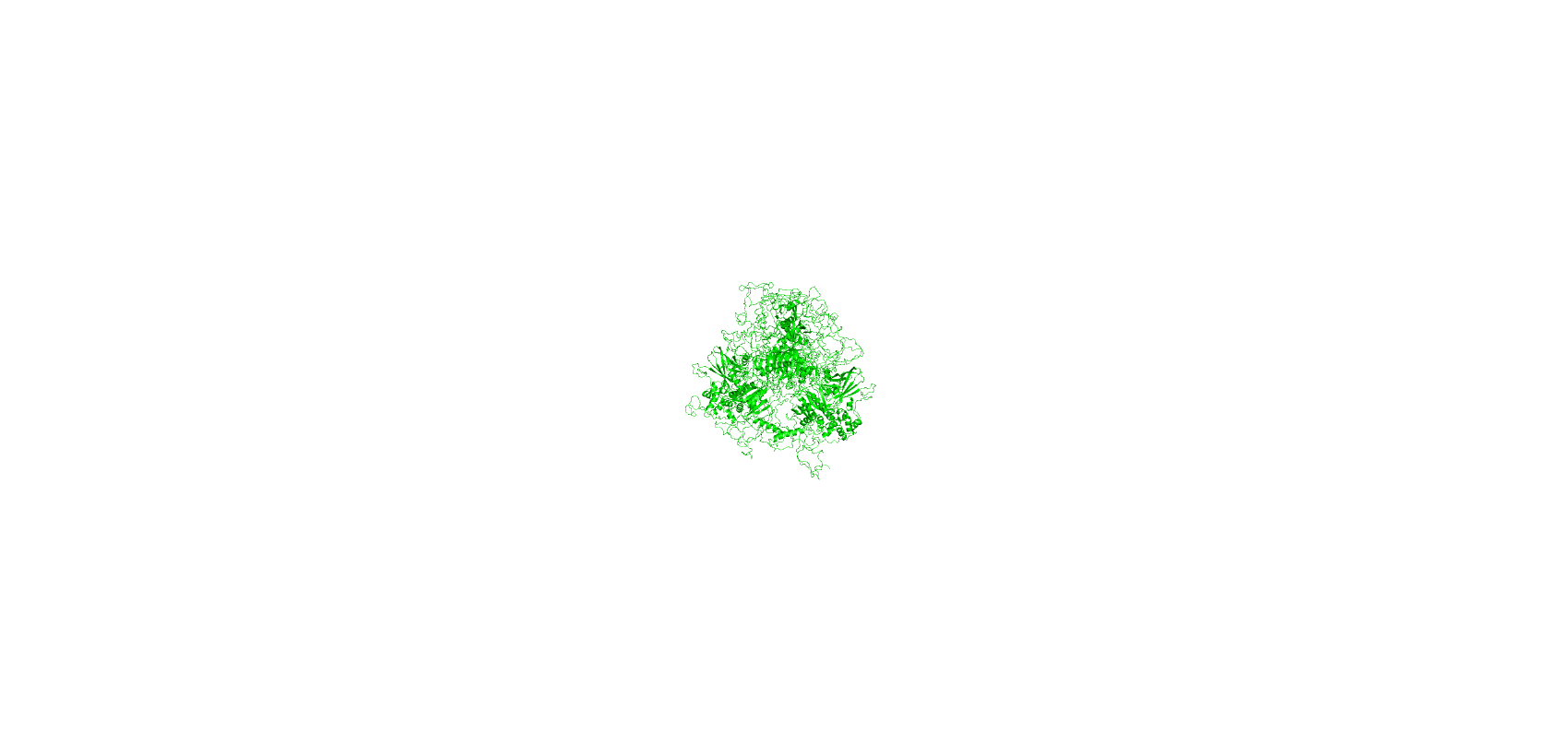

**Fig S10: Bre5-Upb3-Ubiquitin BilboMD refined AF3 model.** The original AF3 model is shown in cyan (χ 2 value of 64.86) and the refined 3-state models (χ 2 value of 2.48) are shown in blue and red and green.

### SEC-SAXS results: Ubp3(407-912) + Ub (Tags removed)

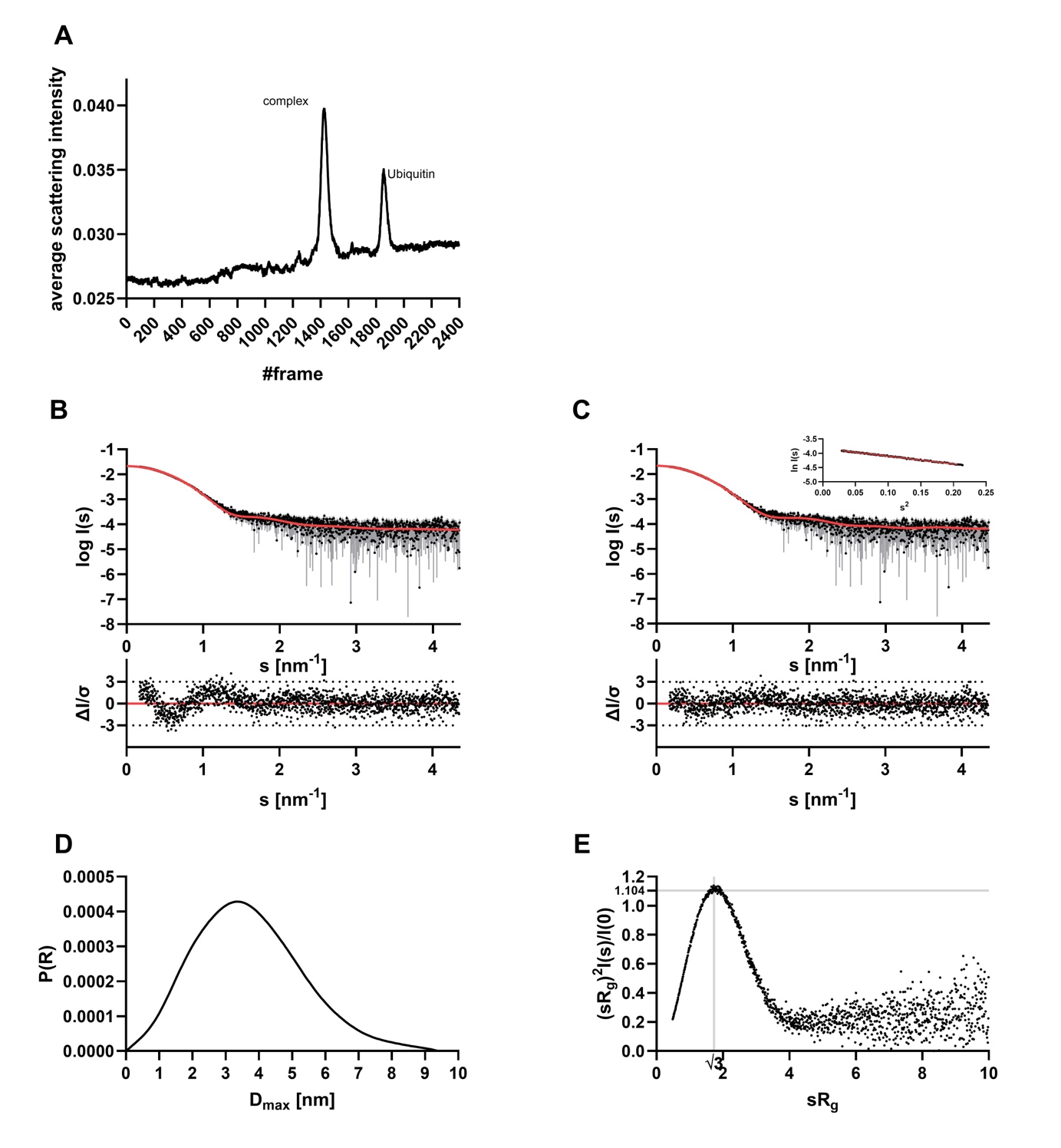

**Figure S11: Small-angle X-ray scattering data of Ubp3-CAT with Ubiquitin. A:** CHROMIXS SEC SAXS elution profile. **B:** Experimental data are shown in black dots, with grey error bars. The theoretical scattering fit of the initial AlphaFold3 model, created with CRYSOL (χ 2 value of 1.376), is shown as red line and below is the residual plot of the data. **C:** Experimental data are shown in black dots, with grey error bars. The best fit model fit (χ 2 value of 1.019) is shown as red line and below is the residual plot of the data. The Guinier plot is added in the right corner. **D:** The Distance distribution function (*p(r)* function) indicate an elongated molecule. **E:** Dimensionless Kratky plot indicate a globular molecule.

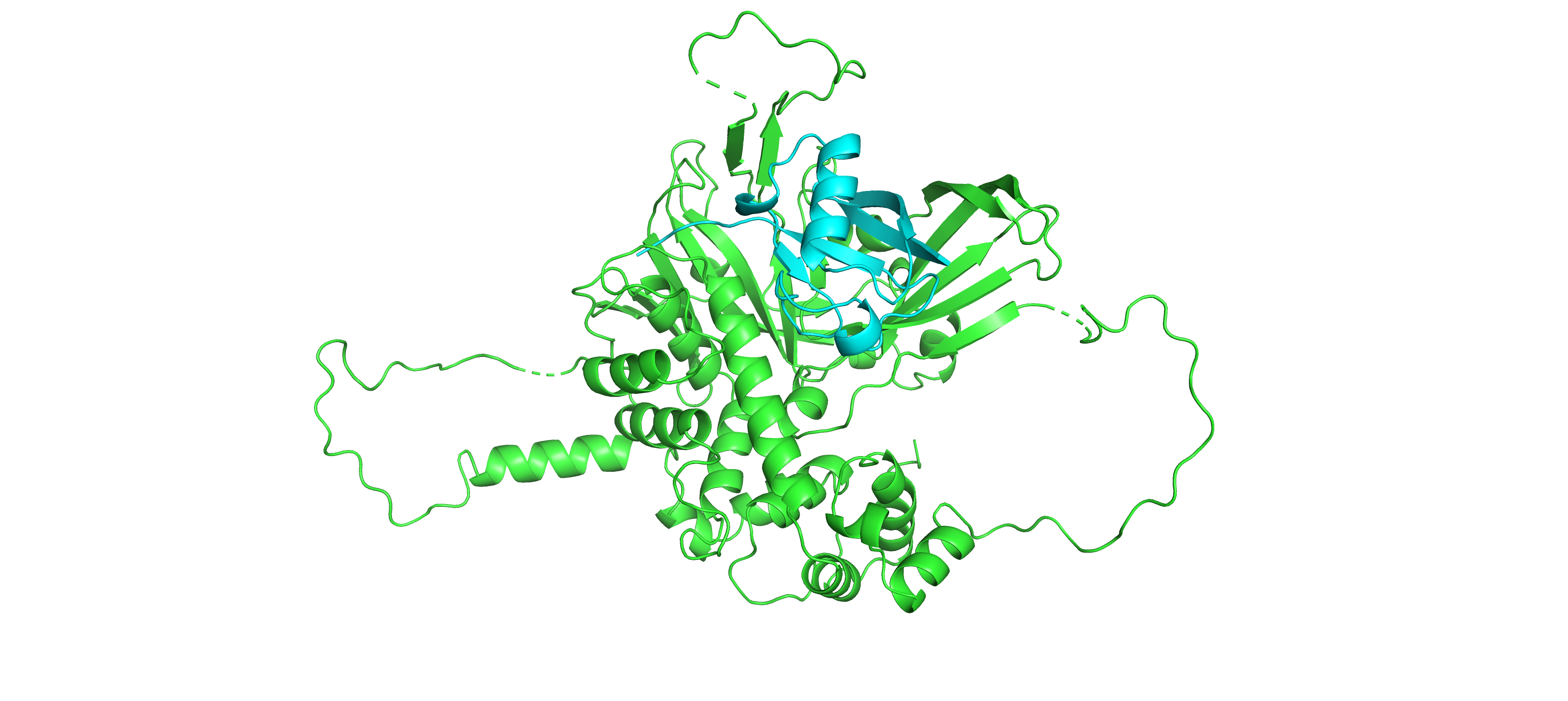

**Figure S12**: Best fit model of Ubp3-CAT with Ubiquitin. Ubp3-CAT is shown in green and Ubiquitin in cyan

### Table S4: Overall SAXS data

| **Data collection parameters** |  | | | | |
| --- | --- | --- | --- | --- | --- |
| SAXS Device | P12, PETRA III, DESY Hamburg [1] | | | BM29, ESRF Grenoble [2] | |
| Detector | PILATUS 6 M | | | PILATUS3 x 2 M | |
| Detector distance (m) | 3.0 | | | 2.827 | |
| Beam size | 120 µm x 200 µm | | | 200 µm x 100 µm | |
| Wavelength (nm) | 0.124 | | | 0.099 | |
| Sample environment | Quartz glass capillary, 1 mm ø | | | Quartz glass capillary, 1 mm ø | |
| Absolute scaling method | Comparison with scattering from pure H_2_O | | | Comparison with scattering from pure H_2_O | |
| Normalization | To transmitted intensity by beam-stop counter | | | To transmitted intensity by beam-stop counter | |
| Scattering intensity scale | Absolute scale, cm^-1^ | | | Absolute scale, cm^-1^ | |
| *s* range (nm^-1^), (s = 4πsin(θ)/λ) | 0.03–7.0 | | | 0.025–5.5 | |
| **Sample** | **Ubp3 catalytic domain** | **Ubp3 CAT with Ubiquitin** | **Bre5-His** | **ALFA-Ubp3-TS** | **Bre5-Ubp3-Ubiquitin complex** |
| Organism | *S. cerivisiae* | *S. cerivisiae* | *S. cerivisiae* | *S. cerivisiae* | *S. cerivisiae* |
| UniProt ID | *Q01477 (407-912)* | *Q01477 (407-912), P0CG63 (1-76)* | *P53741 (1-515)* | *Q01477 (1-912)* | *P53741 (1-515), Q01477 (1-912), P0CG63 (1-76)* |
| Mode of measurement | SEC-SAXS | SEC-SAXS | SEC-SAXS | SEC-SAXS | SEC-SAXS |
| SEC-Column | Superdex 200 increase 10/300 GL | Superdex 200 increase 10/300 GL | Superose6 increase 10/300 GL | Superose6 increase 10/300 GL | Superose6 increase 10/300 GL |
| Flowrate (ml/min) | 0.6 | 0.6 | 0.6 | 0.6 | 0.6 |
| Injection volume (µl) | 100 | 100 | 100 | 100 | 100 |
| Temperature (°C) | 20 | 20 | 20 | 20 | 20 |
| Exposure time (# frames) | 0.995 s (2400 frames) | 0.995 s (2400 frames) | 0.995 s (2400 frames) | 2 s (1200 frames) | 2 s (1200 frames) |
| # frames used for averaging | 36 | 26 | 37 | 17 | 21 |
| Protein buffer | 50 mM Tris pH 8.0, 150 mM NaCl | 50 mM Tris/HCl pH 8.00, 150 mM NaCl, 2% Glycerol | 50 mM Tris pH 8.0, 150 mM NaCl | 20 mM Tris pH 7.5, 500 mM NaCl | 20 mM Tris pH 7.5, 500 mM NaCl |
| Protein concentration (mg/ml) | 12.00 | Ubp3-CAT 5 mg/ml (~90 µM) + 200 µM Ubiquitin | 6.6 | 2.15 | 7.44 |
| **Structural parameters** |  | | | | |
| ***Guinier Analysis (PRIMUS)*** |  | | | | |
| *I*(0) ± σ (cm^-1^) | 0.078 ± 0.00004 | 0.022 ± 0.00004 | 0.0436 ± 0.00014 | 7.332 ± 0.09 | 35.487 ± 0.43 |
| *R*_g_ ± σ (nm) | 2.97 ± 0.008 | 2.85 ± 0.008 | 8.04 ± 0.036 | 4.60 ± 0.09 | 7.75 ± 0.12 |
| *s-range* (nm^-1^) | 0.118 – 0.416 | 0.171 – 0.454 | 0.080 - 0.161 | 0.094 – 0.281 | 0.109 – 0.158 |
| *min < sRg < max limit* | 0.350 – 1.235 | 0.487 – 1.294 | 0.648 – 1.297 | 0.434 – 1.290 | 0.845 – 1.225 |
| Data point range | 1 - 107 | 1 - 102 | 1 - 29 | 1 - 39 | 1 - 10 |
| Linear fit assessment (R^2^) | 0.999 | 0.997 | 0.998 | 0.958 | 0.993 |
| ***PDDF/P(r) Analysis (GNOM 5)*** |  | | | | |
| *I*(0) ± σ (cm^-1^) | 0.078 ± 0.00004 | 0.022 ± 0.00003 | 0.045 ± 0.00014 | 7.747 ± 0.13 | 38.03 ± 0.246 |
| *R*_g_ ± σ (nm) | 3.01 ± 0.003 | 2.86 ± 0.006 | 8.62 ± 0.032 | 5.51 ± 0.17 | 8.72 ± 0.05 |
| *D*_max_ (nm) | 11.11 | 9.39 | 28.89 | 22.03 | 28.28 |
| Porod volume (nm^3^) | 132.40 | 119.77 | 423.06 | 161.53 | 662.81 |
| *s-range* (nm^-1^) | 0.118 – 5.016 | 0.171 – 4.352 | 0.081 – 2.381 | 0.094 – 2.517 | 0.109 – 3.193 |
| χ2 / CorMap P-value | 1.041 / 0.820 | 0.960 / 0.948 | 1.020 / 0.322 | 0.981 / 0.860 | 0.946 / 0.458 |
| **Molecular mass (kDa)** |  | | | | |
| From *I*(0) | n.d. | n.d. | n.d. | n.d. | n.d. |
| From Qp [15] | 66.19 | 66.96 | 265.50 | 115.67 | 414.04 |
| From MoW2 [16] | 58.83 | 65.53 | 147.75 | 115.46 | 347.39 |
| From Vc [17] | 64.72 | 66.02 | 137.15 | 94.70 | 342.67 |
| From Bayesian Inference [18] | 62.35 | 62.35 | 157.05 | 104.62 | 318.45 |
| From Gnnom [19] | 63.40 | 75.90 | 108.80 | 110.00 | 308.20 |
| From sequence | 63.23  (monomer) | 66.33  (1:1 ratio) | 118.66  (dimer) | 107.62 (monomer) | 348.33  (2:2:2 ratio) |
| **Modelling** |  | | | | |
| **CRYSOL (best-fit model)** |  | | | | |
| Template | Model library | Model library | - | - | - |
| Constant subtraction allowed | yes | yes | - | - | - |
| *s-*range for fit (s*_min_* – s*_max_*; nm^-1^) | 0.118 – 5.016 | 0.171 – 4.352 | - | - | - |
| *χ* ^2^, CorMap *P*-value | 1.518 / 3.13e^-09^ | 1.019/ 0.167 | - | - | - |
| **EOM (with default parameters)** |  | | | | |
| Template | - | - | AlphaFold3 | AlphaFold3 | - |
| Constant subtraction allowed | - | - | yes | yes | - |
| *s-*range for fit (s*_min_* – s*_max_*; nm^-1^) | - | - | 0.081 – 2.381 | 0.094 – 2.517 | - |
| *χ* ^2^, CorMap *P*-value | - | - | 1.098 / 0.006 | 0.959 / 0.860 | - |
| **BilboMD (with default parameters)** |  | | | | |
| Template | - | - | - | - | AlphaFold3 |
| *s-*range for fit (s*_min_* – s*_max_*; nm^-1^) | - | - | - | - | 0.109 – 3.193 |
| *χ* ^2^-value | - | - | - | - | 2.48  3-state model |
| **SASBDB accession codes [14]** | - | - | - | - | - |
| **Software** |  | | | | |
| ATSAS Software Version [3] | 3.0.5 /3.2.1 (online server) | | | | |
| Primary data reduction | CHROMIXS [4]/ PRIMUS [5] | | | | |
| Data processing | GNOM [7] | | | | |
| Structure evaluation | CRYSOL [12] | | | | |
| Model library creation | AFflecto [9], RANCH [10, 11] | | | | |
| Flexibility ensemble modelling | EOM [10, 11], BilboMD [13] | | | | |
| Model visualization | PyMOL [20] | | | | |

n.d. =not determined

6.	Guinier, A., *Small-angle X-ray diffraction: application to the study of ultramicroscopic*

*phenomena.* Annales de Physique, 1939. **11**(12): p. 161-237.

20.	PyMOL, *The PyMOL Molecular Graphics System, Version 2.5 Schrödinger, LLC.* 2022.
